# Computational modeling and preclinical validation support targeting somatic instability for Huntington’s disease treatment

**DOI:** 10.64898/2026.01.06.697909

**Authors:** Bryan P. Simpson, Paul T. Ranum, David E. Leib, Luis Tecedor, Ryan C. Giovenco, Icnelia Huerta-Ocampo, Amiel R. Hudley, Noah Smith, Christopher M. Fluta, Christopher P. Cali, Jamie Benoit, Jenna C. Soper, John Connelly, George J. Yohrling, P. Peter Ghoroghchian, Jang-Ho J. Cha, Beverly L. Davidson

## Abstract

Huntington’s disease (HD) is caused by an expanded CAG trinucleotide repeat within the huntingtin (HTT) gene. Genetic modifiers of disease onset and progression in HD implicate somatic instability (SI) of the expanded CAG repeat as a key pathogenic driver, with MSH3 emerging as a leading therapeutic target. Reducing SI, particularly in the most affected neuronal cell type, medium spiny neurons (MSNs) of the striatum, is thus a rational therapeutic strategy for HD. To inform the development of an SI-targeted therapy, we generated a computational model simulating SI in MSNs to infer therapeutic effects on MSN survival resulting from an intervention that reduces SI. The model takes advantage of HD patient data to predict therapeutic benefit across a range of inherited CAG lengths and ages of intervention, considering the degree of target engagement regionally and per cell. To target SI experimentally, we designed an artificial microRNA to lower *MSH3* mRNA (miMSH3) after delivery with AAV-DB-3, a previously described MSN-targeting AAV capsid variant. AAV-DB-3.miMSH3 achieved from 48 to 94% *MSH3* mRNA reduction in MSNs of nonhuman primates (NHPs), which, when modeled, would reduce the composite Unified Huntington Disease Rating Scale change over baseline from 50 to over 120% as well as delay motor symptom onset by many years. AAV-DB-3.miMSH3 also showed robust target engagement in vivo with up to 46% reduction in SI in HdhQ111 mice. The integration of preclinical experimental data and the computational model support the translational potential of AAV-DB-3.miMSH3 as a disease-modifying therapy applicable for HD patients with a broad range of inherited repeat lengths.

**One Sentence Summary:** Predictive modeling to guide therapeutic targeting of disease modifiers in Huntington’s disease.

## INTRODUCTION

Huntington’s disease (HD) is a fatal, autosomal dominant neurodegenerative disease caused by an expanded CAG trinucleotide repeat (>35 CAG) in exon 1 of the huntingtin (*HTT*) gene. The expanded CAG repeat is unstable and undergoes further expansion over a patient’s lifetime through somatic instability (SI) – particularly in vulnerable neuronal populations. In postmortem HD brains, Kennedy and colleagues identified large expansions of up to 1000 CAG repeats in striatal neurons (*1*). Single-cell transcriptomic analyses of HD brain tissue revealed that MSNs with CAG repeat lengths of ≤ 150 maintain normal transcriptional profiles (*2*). In contrast, MSNs with CAG repeat lengths of > 150 display transcriptional dysregulation, a hallmark of HD pathogenesis (*3*). Further, SI is evident in deep layer cortical neurons and in mononuclear cells in blood from HD patients (*4–6*). Together, these findings implicate SI as a major driver of neuronal dysfunction and disease progression, supporting it as a target for disease-modifying therapeutic strategy for HD.

Genome-wide association studies (GWAS) offer critical insights into the role of SI in HD (*7–11*). These studies identified mutS homolog 3 (*MSH3)* variants associated with clinically relevant outcomes, including age at motor onset and rate of disease progression. MSH3, a key component of the mismatch repair (MMR) machinery, binds MSH2 to form the MutSβ complex, which maintains DNA integrity by correcting large insertion/deletion loops that occur during DNA replication (*12*). HD patients with variants associated with reduced *MSH3* expression show slower disease progression. For example, an *MSH3* polymorphism that results in ∼10% lower MSH3 expression reduced SI, slowed disease progression, and delayed disease onset by 1 year (*8*). Likewise, heterozygous loss-of-function mutations in *MSH3* (i.e., 50% MSH3 expression at birth) delayed average age at motor onset by 10.6 years (*13, 14*). These findings provide compelling evidence that even modest reductions in MSH3 expression levels may meaningfully impact the clinical course of HD.

Genetic and pharmacologic studies in mouse and human cell models demonstrate that MSH3 lowering prevents or slows SI, further supporting the link (*14–22*). In MSNs derived from HD patient induced pluripotent stem cells (iPSCs), antisense oligonucleotide (ASO)-mediated knockdown (KD) of MSH3 by 41% proportionally reduced SI while 83% KD roughly halted it (*14*). Notably, MSH3 KD was well tolerated with no evidence of malignant transformation or untoward toxicities (*15, 17–22*). The quantitative relationships between SI lowering, MSN survival, and the associated predicted clinical benefit (e.g., years of delayed clinical progression), however, remain to be defined.

To address this, we designed a computational model to predict the clinical impact of SI-targeted therapies. Foundational work by Handsaker et al. modeled the temporal dynamics of CAG repeat instability at the single MSN level and its relationship to neuronal viability, establishing a framework to quantify SI-driven neuronal loss in HD (*2*). Building on this, we generated a model that integrates natural history data across the spectrum of germline CAG repeat lengths and ages to predict how a SI-lowering therapy impacts neuronal survival and disease stage progression on both a per patient basis and across the HD population. Outcomes are predicated on user-provided information as to the degree of SI reduction and percentage of MSNs impacted, which are information that can be imputed from preclinical studies in mouse and NHP studies. Thus, the model enables facile simulations of outcomes across diverse clinical and interventional metrics, helping to establish clinical translatability thresholds for experimental therapies.

To correlate predicted treatment effects from targeting SI with experimental results, we engineered microRNAs (miRNAs) targeting MSH3 and tested their utility when expressed from AAV-DB-3, a capsid variant that robustly targets MSNs and the vulnerable cortical neuron population in nonhuman primates (NHPs) and mice (*6, 23*). AAV-DB-3.miMSH3 lowered *MSH3* mRNA levels in MSNs throughout the NHP caudate nucleus and putamen and significantly reduced SI in striatal tissues in HdhQ111 mice, a murine model of HD. Cumulatively the data predict that AAV-DB-3.miMSH3 may support neuronal survival and reduce SI expansion in vulnerable HD brain regions, positively impacting the clinical course.

## RESULTS

### Model simulations recapitulate accelerated MSN loss with increasing germline CAG repeat length

To model the clinical benefits from an SI-targeted therapy, we first replicated earlier work simulating CAG repeat expansion in single MSNs (*2*). Our model integrates the previously reported two-phase linear model of somatic CAG repeat expansion rate with a scalable model architecture (*2*). At the base level, individual MSNs are simulated as independent objects with uniquely defined features, including germline CAG repeat length, age, and the potential to be therapeutically modified (Fig. 1A, S1A). As simulated MSNs accumulate somatic CAG repeats over time, they are classified as transcriptionally healthy (<150 CAGs), transcriptionally dysregulated (>150 and <300 CAGs), or dead (>300 CAGs) (Fig. 1B). MSNs modeled with identical starting parameters show expected stochastic variation, following distinct somatic expansion trajectories due to their rates of SI and reflecting the annual probability of variable DNA mismatch repair events, including expansion, contraction, or no change in CAG repeat length (Fig. 1B) (*2*). To represent the collective caudate nucleus and putamen (striatum) in an HD patient, MSN objects are grouped together and assigned to an HD “person” object (Fig. 1C, S1A and B). The simulated HD “person” is configured to age; and, with each simulated year, the impact of CAG repeat expansion on each of the assigned MSN objects is tracked (Fig. 1D, S1C). In the example simulation, the model replicates the stochastic variation in accumulated somatic CAG repeat expansions across MSNs, with some MSNs exhibiting >150 CAGs and the majority remaining in the healthy range at <150 CAGs (Fig. 1D). When the number of MSNs classified as transcriptionally healthy and dysregulated are tracked in a simulated person’s striatum over many decades, the model recapitulates the characteristic late onset and progressive nature of HD after symptom onset. Finally, simulation across a spectrum of CAG repeat lengths and ages explores variation in rates of disease progression at the HD patient population level (Fig. 1E). As an example, with germline CAG repeat lengths of 40, 45, or 50, a shift occurs in the slope of the line, representing fewer healthy MSNs at younger ages with increasing inherited CAG repeat lengths. Thus, the model recapitulates the known inverse correlation between germline CAG repeat length and age of disease onset in HD (Fig. 1F).

**Figure 1:**
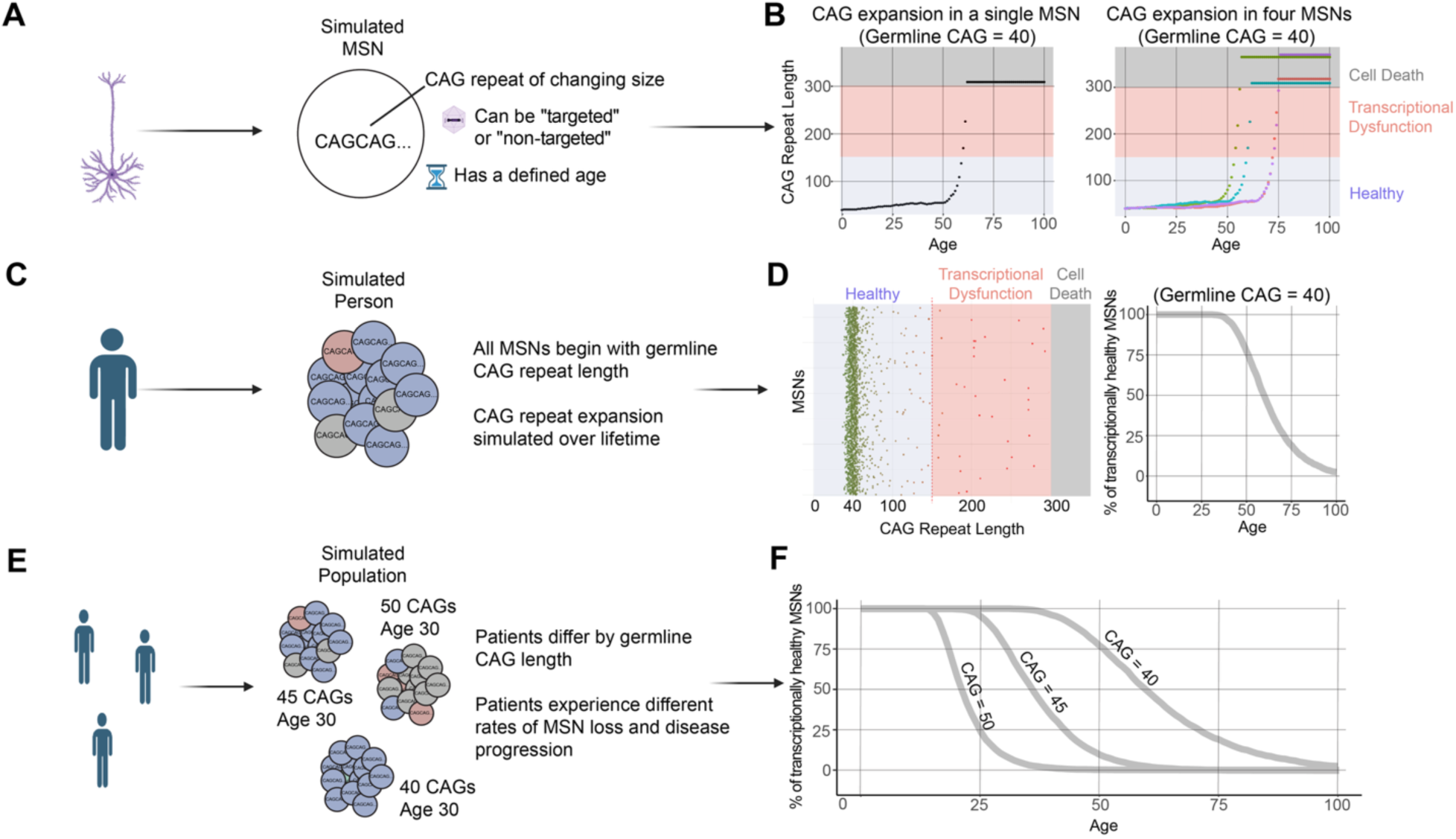
Simulating somatic instability in HD patient MSNs, individuals, and populations. **(A)** Each simulated medium spiny neuron (MSN) contains a defined germline CAG repeat length that can be modified by time and targeting status (targeted or not targeted). (**B)** The left panel represents an example simulation of a single MSN starting with a germline 40 CAG repeat (y-axis) tracked across time from ages 0-100 (x-axis). CAG repeat length is plotted at each simulated age on the y-axis. The right panel depicts the CAG repeat lengths of four simulated MSNs each with different SI rates over time. **(C)** Simultaneous simulation of MSNs across age models disease progression in MSNs of a simulated individual with HD. During simulation, cells are classified as transcriptionally healthy at <150 CAGs (purple), dysregulated at >150 CAGs (red), or dead at >300 CAGs (grey). **(D)** The left panel shows a single frame of the simulation captured at age 40 from an animation of 3,000 MSNs across ages 0-100. The right panel depicts the percentage of healthy MSNs simulated across ages 0-100. The slope of the curve is reflective of the speed of MSN loss and rate of disease progression. **(E)** Simulation of the HD patient population captures the spectrum of patients with a range of germline CAG repeat lengths and a range of ages. **(F)** Three patients with germline CAG lengths of 40, 45, and 50 are simulated and percentages of healthy MSNs are tracked across ages. Panels A, C, and E were created using BioRender.com.

### Simulations of therapeutic targeting of somatic instability predict delayed progression of HD

To simulate the impact of an SI-targeted therapy, the rate of CAG repeat expansion in treated MSNs was modified by two therapeutic parameters: the percentage of targeted MSNs and the percentage of targeted SI reduction (MSH3 KD), using the reported proportional linear relationship between MSH3 levels and SI index (*14, 18*). As a demonstration, therapeutic intervention at age 32 was simulated with 50% MSH3 KD in 50% of MSNs (Fig. 2). These parameters were applied to a simulated individual with an inherited germline CAG repeat length of 40; and, simulations conducted with and without treatment were modeled for healthy and dysregulated MSNs from ages 20 to 80 years old (Fig. 2A-D). [Simulations using the full spectrum of input parameters may be run using our web app (latushdmodel.com)]. By comparing the percentage of remaining MSNs without treatment (Fig. 2C) to with treatment (Fig. 2D), we derive predicted therapeutic benefit; therapeutic benefit is thus defined as the difference in years between onset of a landmark event (defined below) shown as a grey dotted line in Fig. 2E-G in a simulated untreated person versus a simulated treated person (shown by the purple dotted line and purple arrow in Fig. 2E-G). HD clinical landmark events and HD stages were assigned based on longitudinal clinical studies of HD patient cohorts with a variety of germline CAG lengths (*24*). Therapeutic benefit can separately incorporate predicted age of onset for volumetric changes in caudate and putamen (Fig. 2E), the onset of clinical symptoms (motor and cognitive) by HD-ISS stage II (Fig. 2F), and the approximate age of onset for motor symptoms as defined by CAP-100 (Fig. 2G). Finally, we can integrate clinical scores with our model and predict changes over time in simulations of treated and untreated HD patient populations. For example, with treatment at age 32 the rate of cUHDRS (composite Unified Huntington Disease Rating Scale) decline is reduced in the treated condition compared to natural history (untreated), resulting in a steady separation of cUHDRS change over baseline scores (Fig. 2H) (*25*). In summary, simulating the effect of an SI-targeted therapy over a patient’s lifetime enables characterization of therapeutic benefit as years of delay in onset of several HD landmarks and reduced decline in cUHDRS.

**Figure 2:**
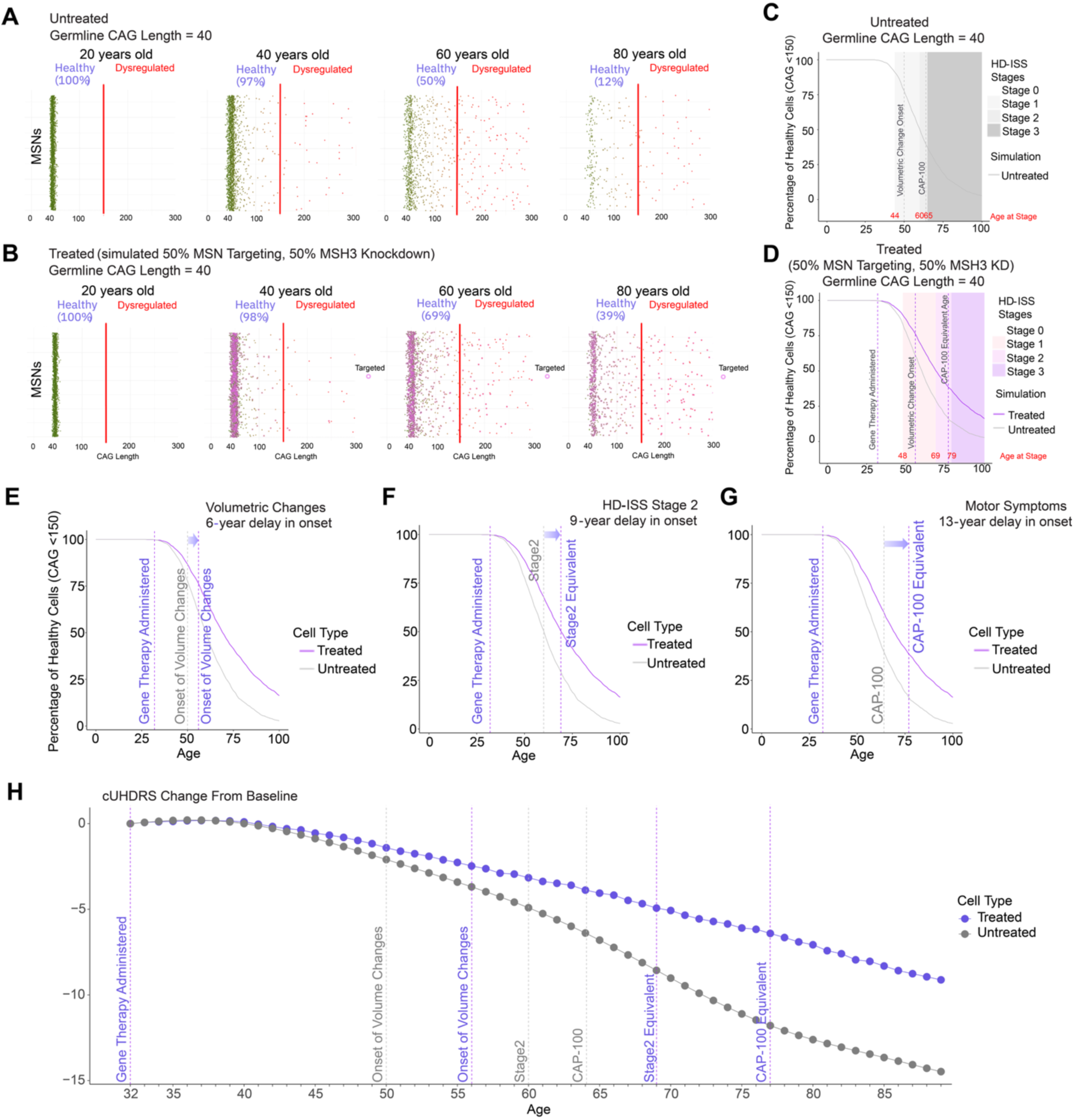
Simulating MSH3 knockdown on MSN survival over a patient lifetime. **(A-B)** Still frames from an animation of untreated **(A)** and treated **(B)** HD patients are shown from example simulations of 3000 MSNs that depict somatic CAG repeat expansion in individual MSNs at 20-year intervals from 1-100 years. **(A)** Each point represents an untreated MSN and simulated CAG length (x-axis). The red line represents the 150 CAG threshold for transcriptionally healthy MSNs (<150 CAG; green) and dysregulated unhealthy MSNs (>150 CAG; red) progressing toward cell death (>300 CAGs). **(B)** Simulated MSNs are shown with SI-targeted therapy administration at 32 years old and 50% of MSNs targeted with 50% MSH3 knockdown. Targeted MSNs are circled in pink. **(C)** The percentage of untreated healthy MSNs (<150 CAGs) are tracked and plotted across a simulated lifetime (0-100 years). Shaded background panels represent the indicated HD-ISS stage, age (shown in red) when a HD-ISS stage is reached, and dotted lines represent a HD milestone event (measurable volumetric change and CAP-100 equivalent age). **(D)** A treated purple line is compared to the untreated grey line from **(C)** and tracks the percentage of healthy treated MSNs after administration at 32 years old. **(E-G)** The model tracks therapeutic benefit by determining the age and healthy MSN number coinciding with a HD landmark event (onset of striatum volume change, motor symptoms, or HD-ISS stage). **(H)** The model enables prediction of cUHDRS change from baseline between simulated treated (purple dotted line) and untreated Enroll-HD natural history cohorts (grey dotted line).

### Age of intervention impacts clinical benefit

Next, we predicted therapeutic benefit for individuals inheriting from 37 to 60 germline CAG repeat lengths (*26*) with various ages of intervention (5 to 60 years of age) (Fig. 3A-C). Because a therapeutic benefit of 5 years is a reasonable threshold for a SI-targeted therapy to show measurable and meaningful efficacy in an HD patient clinical trial, this was defined as our minimal therapeutic threshold. We then simulated outcomes based on percentage of MSNs targeted and degree of MSH3 KD. With only 25% KD in 25% of MSNs, the model predicts five years of therapeutic benefit for HD patients with low germline CAG repeat lengths that undergo treatment early in life (Fig. 3A). We next simulated MSH3 KD levels of 50%, 75%, and 100%, and held the percentage of targeted MSNs at 50% (Fig. 3B), or, simulated variable MSN targeting rates of 50%, 75%, and 100%, and kept MSH3 KD levels at 50% (Fig. 3C). The output reveals that neuron survival is maximized by early intervention, but also that the level of MSH3 reduction is more impactful than reducing modest amounts of MSH3 in all cells. Thus, MSNs targeted by an SI-targeting therapy will still experience rapid CAG repeat expansion if the level of MSH3 KD is minimal or if the targeted MSNs already have advanced CAG repeat lengths. Conversely, if SI is prevented in a low number of MSNs through robust MSH3 KD, those MSNs could indefinitely persist at a healthy CAG repeat length. Collectively, these simulations bracket 50% MSN targeting with 50% MSH3 KD as targets for therapeutic impact for a large cross section of the HD patient population (as defined by a range of CAG repeat lengths and ages at intervention).

**Figure 3:**
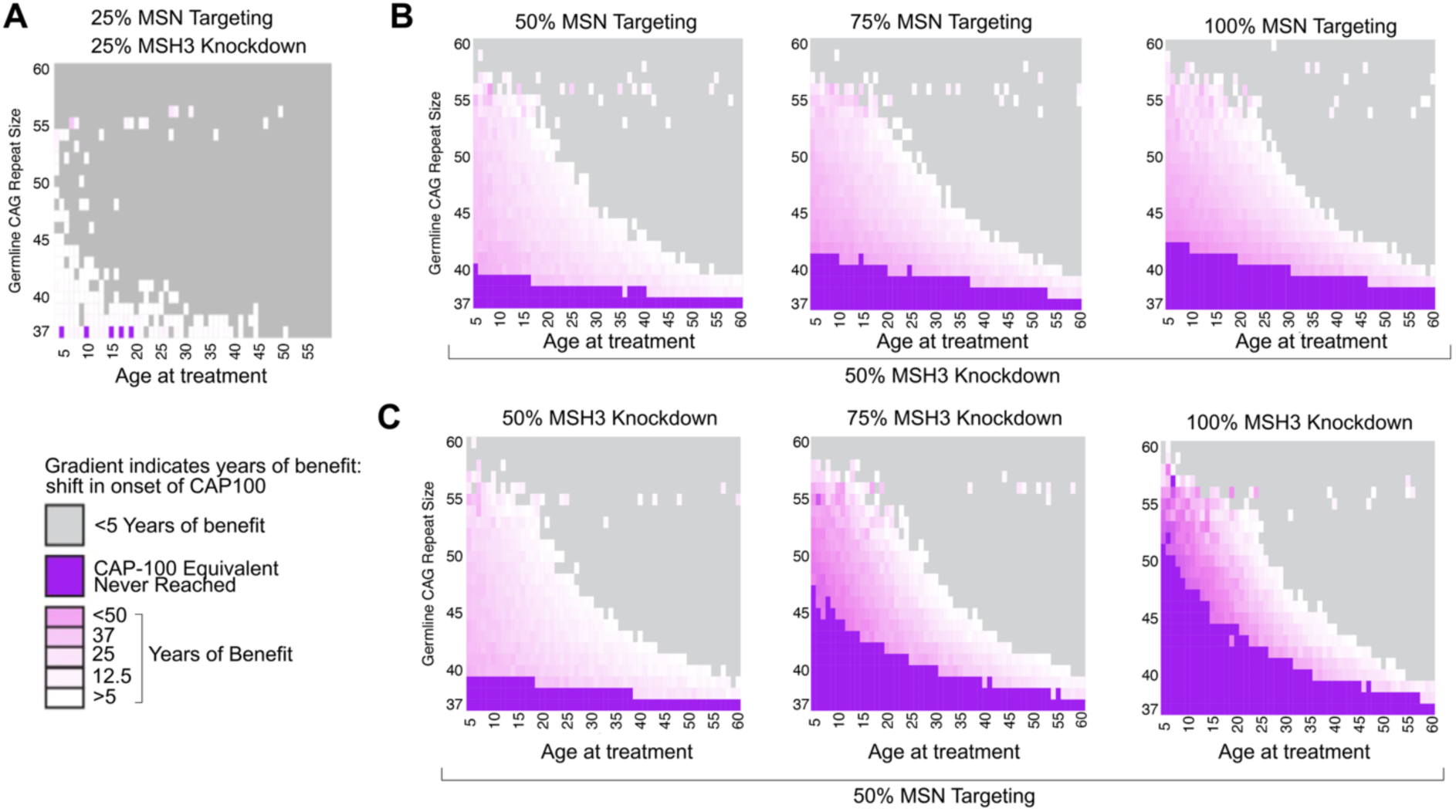
Predicting therapeutic benefit across a spectrum of simulated HD patients. **(A-C)** Heatmaps depicting the simulated therapeutic benefit for a SI-lowering therapy in a range of treated patients with germline CAG repeat lengths from 37-60 CAGs (y-axis) and ages at treatment from 5-60 years of age (x-axis). The degrees of predicted therapeutic benefit as measured by years of delay to CAP-100 equivalent are depicted in the heatmaps (shown in the heatmap key; less than 5 years of benefit shown in grey, CAP-100 equivalent never reached shown in purple, and benefit between >5 and <50 years shown by the white to pink gradient). **(A)** Heatmap depicting the simulated range of therapeutic benefit achieved by targeting 25% of MSNs with 25% MSH3 knockdown per MSN. **(B)** Heatmaps depicting the range of therapeutic benefit with 50% MSH3 knockdown per MSN and a range of MSN targeting percentages (50%, 75%, and 100%). **(C)** Heatmaps depicting the range of therapeutic benefit with targeting 50% of MSNs and a range of MSH3 knockdown percentages (50%, 75%, and 100%) per MSN.

### Artificial miRNAs potently reduce MSH3 expression in cells and wildtype mice

Artificial miRNAs were designed to target both NHP and human *MSH3* mRNA (miMSH3). *In silico* predictions were done with siSPOTR (*27*) - a bioinformatic tool for design of highly specific and potent miRNA sequences. Two candidates, miMSH3-06 and miMSH3-11, emerged from the *in vitro* screening for potency and strand bias, showing minimal passenger strand loading and potent reduction of endogenous human and mouse *MSH3* mRNA in HEK293 cells and N2a cells, respectively (Fig. S2A-E).

For *in vivo* testing, miRNAs were packaged in the capsid variant, AAV-DB-3 (*23*). The miMSH3-06 and miMSH3-11 guide sequences were tested in the miR-30 scaffold (AAV-DB-3.miR30-miMSH3-06, AAV-DB-3.miR30-miMSH3-11) and in the miR-451 scaffold (AAV-DB-3.miR451-miMSH3-06, AAV-DB-3.miR451-miMSH3-11). miMSH3-06 and miMSH3-11 embedded in the miR-30 scaffold outperformed the same miMSH3s embedded in the miR-451 scaffold, demonstrating greater *Msh3* mRNA KD in the striatum of WT mice after intrastriatal delivery (Fig. S3A-C). Finally, miMSH3-06 and miMSH3-11 demonstrated efficient miRNA processing with high 5’-end accuracy (>90%) and favorable guide-to-passenger strand ratios in HEK293 cells (>9:1) and WT mouse striatal tissues (>85:1) (Fig. S4A-B).

As miRNA sequences can be partially complementary to unintended mRNAs, resulting in off-target gene expression effects, the off-target profiles of miMSH3-06 and miMSH3-11 were evaluated. Both miMSH3-06 and miMSH3-11 showed low off-target KD activity and robust KD of *MSH3* mRNA in HEK293 cells (Fig. S5). miMSH3-06 showed fewer off-target hits by low fold change (<1 log2 fold change) (Fig. S5). Overall, these results supported the selection of miMSH3-06, hereafter called miMSH3, as the lead miRNA candidate for further testing in NHPs and in HdhQ111 knock-in (KI) mice.

### AAV-DB-3.miMSH3 reduces MSH3 mRNA and protein levels in NHP caudate and putamen

A four-week in-life dose range finding (DRF) study was performed in rhesus macaques to identify effective doses of AAV-DB-3.miMSH3 for target reduction and percent MSN transduction after intraparenchymal (IPa) globus pallidus (GP) infusion (Fig. 4A; Table S1). Bulk tissue analyses showed dose-dependent increases in AAV-DB-3 biodistribution (Fig. 4B) and miMSH3 expression (Fig. 4C) in the GP, caudate, and putamen. AAV-DB-3 biodistribution was evident throughout brain regions, and as expected, there was limited detection in peripheral tissues (Fig. S6). In bulk GP tissue, *MSH3* mRNA levels were reduced by 31%, 52%, and 47% relative to vehicle control for the low-, mid-, and high-dose groups, respectively (Fig. 4D). These reductions in mRNA corresponded to 22%, 47%, and 34% reductions in MSH3 protein levels, respectively (Fig. 4E). In the caudate, *MSH3* mRNA levels were reduced by 14%, 16%, and 39%, and MSH3 protein levels were reduced by 12%, 23%, and 28% at the low, mid, and high doses, respectively (Fig. 4D-E). Finally, in the putamen, *MSH3* mRNA levels were reduced by 9%, 16%, and 24% at the low, mid, and high doses, respectively. MSH3 protein levels were unchanged at the low dose and reduced by 19% at both the mid and high doses (Fig. 4D-E). Together, MSH3 mRNA and protein were generally reduced in bulk tissue in a dose-dependent manner in the caudate and putamen. Additionally, miMSH3 processing was favorable in the putamen of NHPs with >99% 5’-end accuracy and >30:1 guide-to-passenger strand ratio in the NHP putamen (Fig. S4A-B).

**Figure 4.**
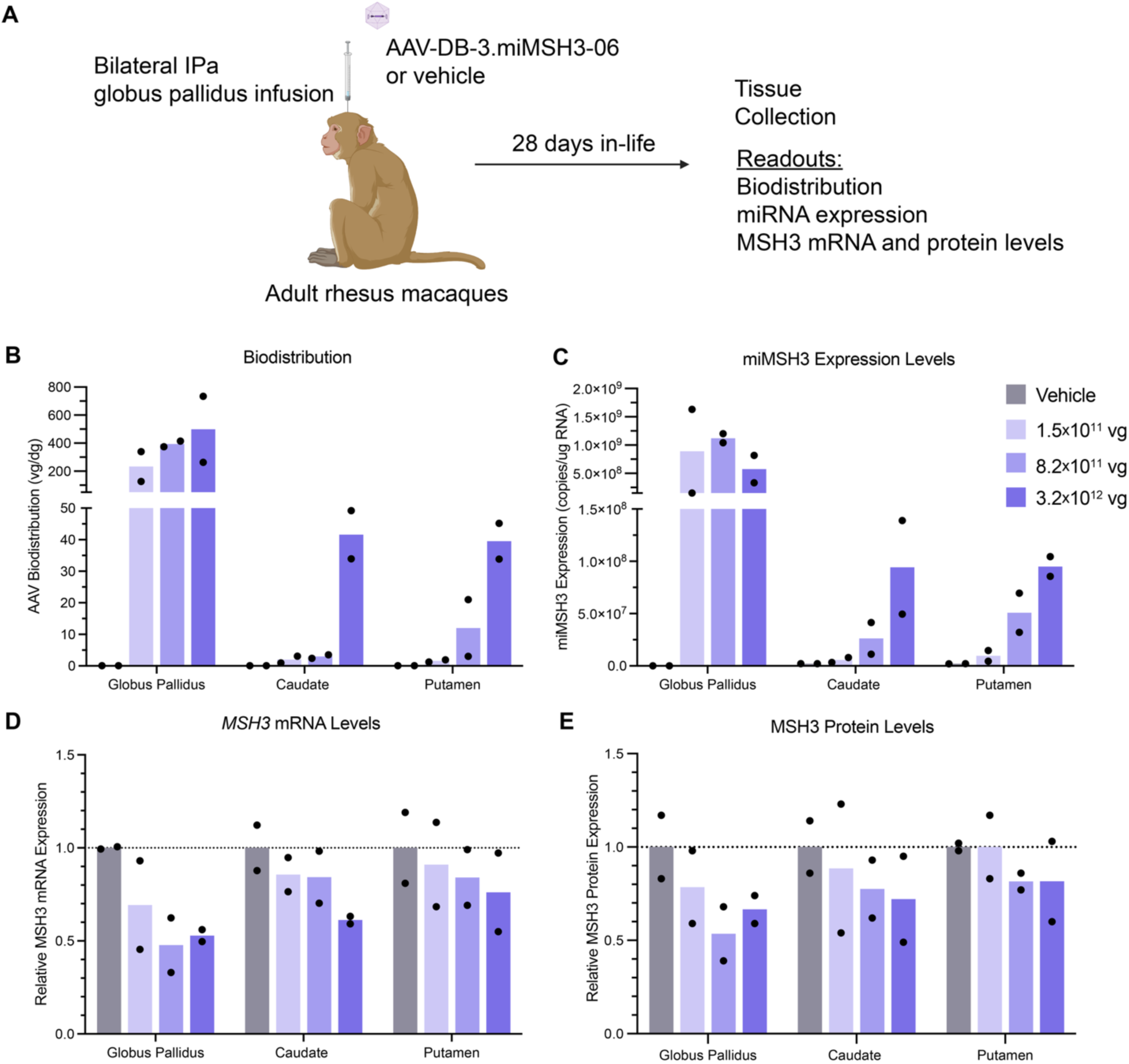
Target engagement and MSH3 lowering in NHP brain. **(A)** Study design for the nonhuman primate dose range finding study in rhesus macaques. **(B)** AAV vector biodistribution (vector genomes per diploid genome) in the globus pallidus (GP), caudate nucleus, and putamen of vehicle and treated rhesus macaques by droplet digital PCR (ddPCR) after bilateral intraparenchymal (IPa) GP administration of AAV-DB-3.miMSH3-06. **(C)** miMSH3 expression levels normalized to copies per µg of RNA by stem-loop RT-qPCR. **(D)** *MSH3* mRNA levels normalized to *TBP* and relative to vehicle control by RT-qPCR. **(E)** MSH3 protein levels relative to vehicle control by Jess capillary-electrophoresis immunoassay. Data are shown as group mean (N=2) with data points from each animal. Panel A was created using BioRender.com.

The bulk tissue sampling confirmed biological activity of AAV-DB-3.miMSH3 and target engagement but likely underestimates MSH3 KD in MSNs as non-transduced cells (e.g. astrocytes, oligodendrocytes, etc.) dilute the MSN-specific KD effect due to AAV-DB-3’s tropism (*23*). Indeed, MSNs, while accounting for 85% of neurons in the striatum, are only 34.4% of the overall cell population (Fig. S7) (*28*).

To quantify *MSH3* mRNA levels specifically in *PPP1R1B^+^* (DARPP-32^+^) MSNs in NHP caudate and putamen, single-cell RNAscope fluorescence *in situ* hybridization (FISH) was performed. An automated counting algorithm was used to quantify the puncta in the cytoplasmic compartment, affording quantification of *MSH3* target engagement and KD (Fig. 5A-B). We prioritized the cytoplasmic signal because miRNAs primarily target mRNAs in the cytoplasm (*29*). Quantitative analysis of confocal images from RNAscope experiments showed dose-dependent decreases in cytoplasmic *MSH3* mRNA in all MSNs profiled in the striatum treated with AAV-DB-3.miMSH3 (Fig. 5C). Compared to vehicle-treated animals, cytoplasmic *MSH3* mRNA levels in striatal MSNs were reduced by 48%, 61%, and 94% in the low-, mid- and high-dose groups, respectively (Fig. 5C). Lowering of *MSH3* mRNA was also observed in MSN nuclei, suggesting a fraction of nuclear *MSH3* mRNA was accessible to RNA interference (RNAi) KD or that KD occurred indirectly in the nuclear compartment, whereby both compartments reached an equilibrium (Fig. S8). Together, these results demonstrate robust reduction of *MSH3* mRNA in NHP striatal neurons, exceeding our predicted therapeutic threshold for successful translation of an SI-targeted therapy.

**Figure 5.**
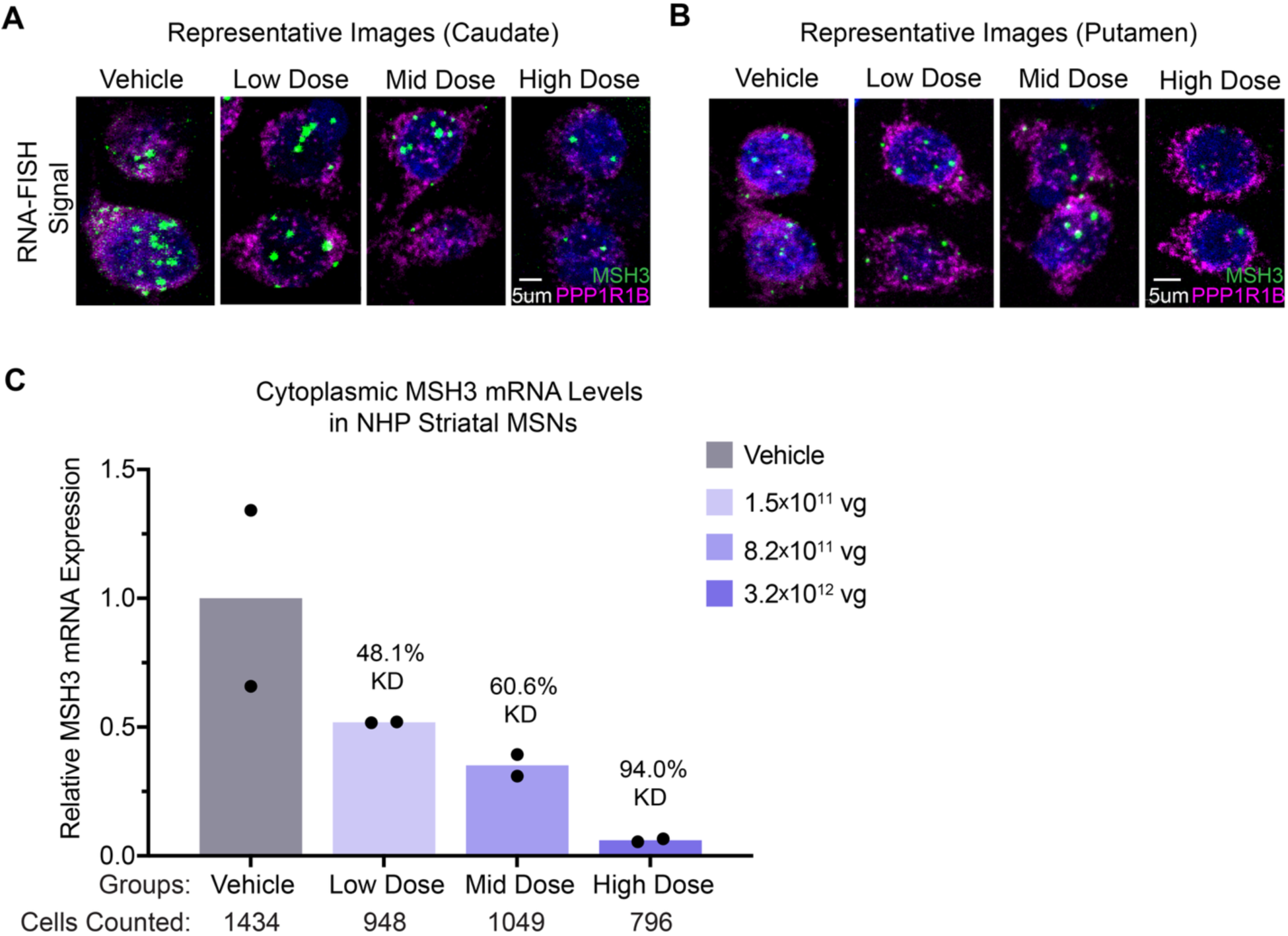
Dose-dependent lowering of *MSH3* mRNA in striatal MSNs. **(A-B)** Representative RNAscope FISH confocal microscopy images of *MSH3* mRNA puncta (green) in *PPP1R1B+* MSNs (magenta) in the caudate **(A)** and putamen **(B)** of NHPs. Hoechst stain (blue) was used to counterstain nuclei. **(C)** Cytoplasmic *MSH3* mRNA puncta count in *PPP1R1B+* MSNs of the striatum (caudate and putamen) were determined with an automated cell segmentation and puncta counting algorithm. The percentages of cytoplasmic MSH3 mRNA knockdown (KD) levels in all MSNs profiled in the caudate and putamen relative to vehicle control are shown above each treatment group. Total MSN counts are displayed below each group. Data are shown as group mean (N=2) with data points from each animal.

### NHP data predict a measurable benefit in HD

Next, the RNAscope data on MSH3 mRNA lowering was incorporated into the computational model to simulate the translational performance of the low, mid, and high dose treatments of AAV-DB-3.miMSH3. We set the *MSH3* mRNA KD level in MSNs for each dose tested in NHPs across a range of inherited CAG repeat lengths and ages of intervention (20-60 years old) (Fig. 6A-D). The range of 40-50 captures most germline CAG repeat lengths present in the HD patient population (shown by the red-yellow color gradient CAG repeat length incidence indicator) (*26, 27*). We again applied a minimum threshold of 5 years as therapeutically relevant to facilitate comparison. The range of patients meeting the 5-year therapeutic threshold minimum are denoted as black outlined rectangles; for reference, they are shown on the low, mid, and high dose heatmaps (Fig. 6A-D). Simulation of the *MSH3* mRNA KD levels of 48.1%, 60.6%, and 94.0% in MSNs profiled at the low, mid, and high doses, respectively, demonstrated that all doses exceeded the minimum therapeutic threshold, supporting the decision to advance AAV-DB-3.miMSH3 (Fig. 6B-D). Remarkably, all doses also showed the potential to prevent a subset of patients from ever developing motor symptoms (>50 years of delay in onset to CAP-100 equivalent score; Fig. 6B-D, dark purple rectangles). The predicted therapeutic effect of MSH3 KD is further demonstrated by a simulated reduction in cUHDRS change over baseline by 53%, 40%, and 126% relative to natural history in the low, mid, and high doses, respectively, at three years post-treatment (Fig. 6E). Overall, modeling suggests that the observed *MSH3* mRNA KD levels in NHP MSNs may translate to substantial clinical benefit across a range of HD patients.

**Figure 6.**
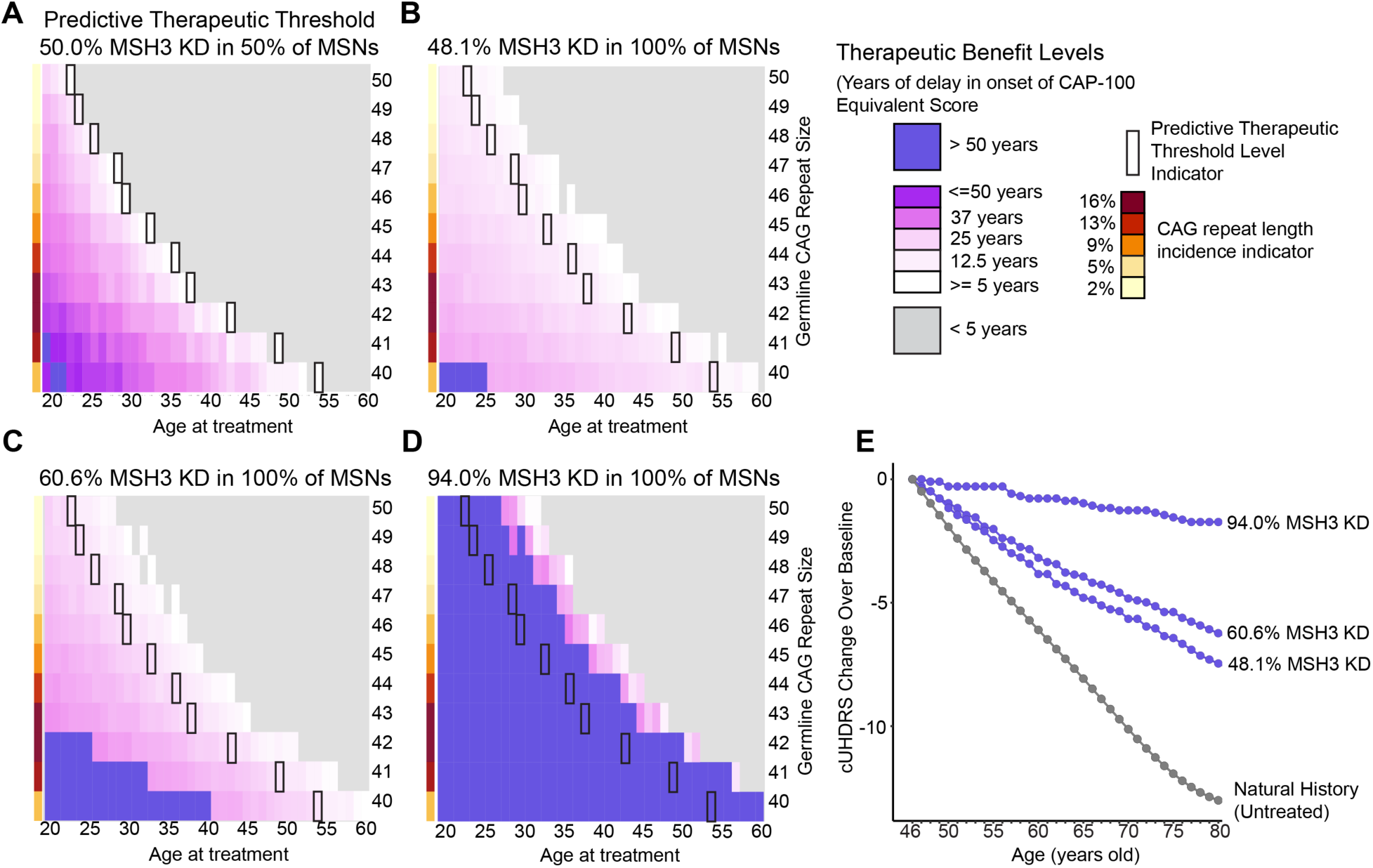
Clinical therapeutic benefit predicted at all doses tested in NHPs. **(A-D)** Heatmaps depicting the simulated range of therapeutic benefit for patients treated with an SI-lowering AAV gene therapy over a lifetime of 100 years. Each heatmap represents a range of patients with germline CAG lengths from 40-50 CAG repeats (y-axis) and age at treatment (x-axis). The degrees of predicted therapeutic benefit as measured by years of delay to CAP-100 equivalent are depicted in the heatmaps (shown in the heatmap key; less than 5 years of benefit shown in grey, CAP-100 equivalent was never reached shown in purple, and benefit between >5 and <50 years shown by the gradient from white to pink). **(A)** The range of therapeutic benefit achieved after simulating 50% of MSNs transduced and 50% MSH3 KD per transduced MSN. This threshold is denoted by black bordered heatmap cells in all four heatmaps. **(B-D)** Heatmaps depict the predicted range of therapeutic benefit for HD patients treated with the NHP DRF dose equivalents of AAV-DB-3.miMSH3 at the low dose (1.5×10^11^ vg) **(B)**, mid dose (8.2×10^11^ vg) **(C)**, and high dose (3.2×10^12^ vg) **(D)** based on the RNAscope FISH *MSH3* mRNA KD levels detected in the NHP caudate and putamen (Figure 5C). CAG repeat length incidence is shown by the red to yellow color gradient column (*42*). **(E)** To characterize predicted therapeutic performance using cUHDRS across the observed percentages of MSH3 knockdown, we simulated these therapeutic performance levels in a hypothetical patient with 42 CAGs treated at the age of 46.

### AAV-DB-3.miMSH3 reduces somatic instability

The effects of AAV-DB-3.miMSH3 on SI were next assessed in the HdhQ111 KI mouse model of HD (Fig. 7A). The HdhQ111 KI mouse exhibits robust somatic CAG expansion in the striatum, similar to observations in post-mortem human HD brains (*1, 30*). Study groups included untreated and vehicle-treated heterozygous HdhQ111 KI mice, as well as heterozygous HdhQ111 KI mice dosed with AAV-DB-3.miMSH3 at 5×10^9^, 1.5×10^10^, 5×10^10^ or 1.5×10^11^ vg per mouse via bilateral intrastriatal injections. Mice were injected at 8 weeks of age and euthanized 16 weeks later to assess vector biodistribution, target KD and SI.

**Figure 7.**
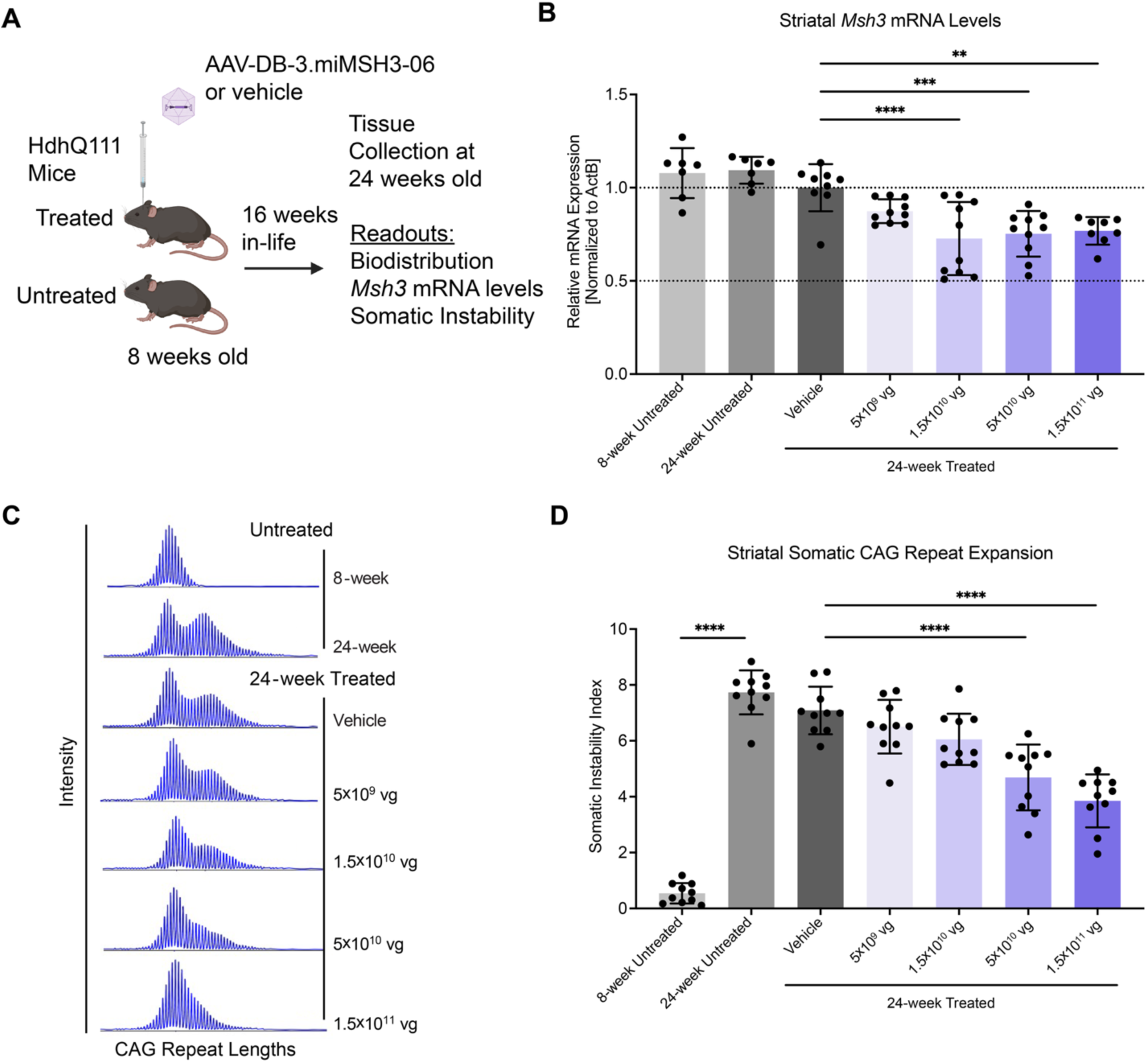
Dose-dependent slowing of somatic CAG repeat instability in HdhQ111 mice. **(A)** Study design for the pharmacology study in heterozygous HdhQ111 KI mice. **(B)** *Msh3* mRNA levels in the striatum of untreated and treated HdhQ111 mice by RT-qPCR. *Msh3* mRNA levels were normalized to *ActB* mRNA and shown as the group mean relative to vehicle control (N=7, 8 week untreated; N=7, 24 week untreated; N=9, vehicle; N=10, 5×10^9^ vg; N=10, 1.5×10^10^ vg; N=10, 5×10^10^ vg; N=8, 1.5×10^11^ vg; one-way ANOVA with Dunnett’s post hoc analysis, \*\**p* < 0.01, \*\*\**p* < 0.001, \*\*\*\**p* < 0.0001). **(C)** Fragment analysis traces shown in blue depict the intensity of CAG repeat length alleles in untreated, vehicle and AAV-DB-3.miMSH3-06 treated mice by PCR and capillary electrophoresis. **(D)** Somatic instability index in untreated and treated mice (N=10/group; one-way ANOVA with Dunnett’s post hoc analysis, \*\*\*\**p* < 0.0001). Data are shown as the mean and error bars represent SD. The percent reduction in SI relative to vehicle control is shown above each treatment group. Panel A was created using BioRender.com.

In bulk striatal tissue, AAV-DB-3 biodistribution was dose-dependent and highly specific to the brain with minimal detection in peripheral tissues (Fig. S9A). Striatal *Msh3* mRNA levels were reduced by 13%, 27%, 25%, and 23% at the 5×10^9^, 1.5×10^10^, 5×10^10^ and 1.5×10^11^ vg/mouse doses, respectively, compared to 24-week-old vehicle-treated controls (Fig. 7B). Furthermore, striatal *Msh3* mRNA levels in the 1.5×10^10^, 5×10^10^ and 1.5×10^11^ vg/mouse dose groups were significantly reduced compared to the 24-week-old vehicle-treated controls (Fig. 7B). These data suggest that increasing levels of striatal biodistribution do not correspond to a clear dose-response in *Msh3* lowering, which appeared saturated above the 1.5×10^10^ vg/mouse dose level.

To assess the impact of AAV-DB-3.miMSH3, SI was measured in HdhQ111 KI mice. SI significantly increased from 8 to 24 weeks of age in untreated control striata; and, vehicle-treated 24-week-old controls did not differ significantly from 24-week-old untreated controls (Fig. 7C-D). AAV-DB-3.miMSH3 reduced the SI index in the striatum by 8%, 15%, 34%, and 46% at doses of 5×10^9^, 1.5×10^10^, 5×10^10^ and1.5×10^11^ vg/mouse, respectively, compared to vehicle-treated controls (Fig. 7C-D). Both the 5×10^10^ and 1.5×10^11^ vg/mouse dose levels significantly reduced SI relative to vehicle controls (Fig. 7D). In an independent validation study using miMSH3-11, intrastriatal delivery of AAV-DB-3.miMSH3-11 in HdhQ111 mice at a single dose of 5×10^10^ vg/mouse showed significant lowering of striatal *Msh3* mRNA by 20% and a concomitant significant reduction in striatal SI index by 43% (Fig. S10A-E). Overall, AAV-DB-3.miMSH3 reduced *Msh3* in HdhQ111 KI mice, supporting target engagement and effective reduction of SI.

## DISCUSSION

SI of the expanded CAG repeat has emerged as a driver of HD pathogenesis. Genetic studies from HD patient cohorts strongly implicate SI as the determinate of the rate of disease onset and progression (*5, 6*, *7*, *8*). The potential clinical benefit of SI-targeting therapies – particularly when and in whom they should be applied – remains uncertain. As the HD therapeutic landscape evolves toward developing DNA repair-modulating strategies, there is a need for translational frameworks that quantitatively connect target engagement and impact on SI to predicted clinical outcomes. Our work addresses this gap by integrating computational modeling along with patient natural history and preclinical data to predict the potential clinical benefits of targeting SI with an MSH3-lowering intervention. It further helps to establish predicting therapeutic thresholds and to identify patient populations most likely to benefit.

We build on previous work that has elucidated the correlation between CAG repeat expansion rates and neuronal viability to now assess the impact of SI-modifying therapies on overall clinical progression over a range of different interventional and patient characteristics. This is accomplished by simulating the lifetime impact from inheriting a given CAG length and its associated impact on MSNs in ‘individuals,’ in the presence and absence of SI modifying therapies. We applied the model across a broad HD patient population with variable germline CAG lengths and ages (*2*). By varying MSH3 KD levels, MSN coverage, treatment age, and inherited CAG repeat length, the model predicts that partial suppression of SI preserves healthy neurons. This outcome is validated by human GWAS data and mouse studies, showing benefit from loss of MSH3 function (*7–10, 20, 21*). Notably, our model is broadly applicable to any SI-lowering approach and independent of modality (e.g., antisense oligonucleotides, RNAi, gene editing, or small molecules), as it defines the quantitative relationship between the degree of SI suppression in targeted MSNs and the predicted benefits on disease outcomes. Thus, we have established a scalable and modular platform for evaluating SI-targeted therapies, enabling investigators to assess hypothetical interventions against predefined efficacy thresholds and to validate them in preclinical models. Importantly, even modest therapeutic performance (50% MSN targeting and 50% MSH3 knockdown) is predicted to delay motor onset by ≥ 5 years for a wide range of HD patients, demonstrating that potentially measurable and meaningful benefits may be achievable even without complete target suppression. Finally, the model simulations of targeting SI, as well as data from human HD patients and animal models, underscores the importance of lowering MSH3 early; this would have the most impact on modifying the disease course before the CAG repeats expand beyond a pathogenic threshold in a significant number of MSNs, preserving their longevity (*2, 4, 22*). The model also supports that treating later in the disease course, and with sufficient target knockdown, can provide meaningful benefits, albeit to a lesser degree.

We previously engineered the AAV-DB-3 capsid variant for efficient and selective transduction of MSNs within the basal ganglia and deep layer cortical neurons (*23*). The work presented here integrates the MSN-tropic properties of AAV-DB-3 with a genetically validated SI-modifying target, MSH3, and our predictive model to guide preclinical development. The computational model identified the levels of MSH3 reduction required to yield clinical benefit, which we show can be achieved even at modest dosing in NHPs. Complimentary HdhQ111 mouse studies demonstrated dose-dependent slowing of SI consistent with the predicted mechanism of action. Together, the convergence of empirical and simulated data illustrates how predictive translational modeling and preclinical testing can inform a therapeutic approach, enabling the establishment of efficacy thresholds, dose selection, and patient population criteria.

If effectively translated, SI-targeting interventions such as MSH3 KD could expand the therapeutic options beyond symptomatic HD patients to include presymptomatic *HTT* mutation carriers. In this population, slowing or stopping SI before the onset of SI-driven transcriptional dysregulation could preserve neuronal integrity and prevent disease. Moreover, early intervention according to the model-informed thresholds could enable preventative gene therapy paradigms. Because SI occurs on the mutant allele, MSH3 lowering provides an indirect allele-selective strategy that could limit the expression of the toxic, hyperexpanded mHTT species, including HTT1a, and the formation of aggregates (*21, 31*).Additionally, SI targeting may lessen the need for HTT-lowering approaches, though combined strategies may yield greater benefit. Dual-targeting of MSH3 and mHTT – or additional targets such as PIAS1 – could address complementary and independent mechanisms of neuronal toxicity (*21, 32*). Beyond MSH3, the model provides a generalizable framework for evaluating other SI-modulating targets (e.g., PMS1, FAN1) and combinatorial strategies targeting other DNA repair pathways (*20, 33, 34*). This modularity allows for expansion to assess additive or synergistic benefits from multimodal SI-targeting therapies.

More broadly, our computational and translational modeling framework could be adapted to other repeat expansion disorders where SI contributes to disease progression. However, data describing the rate of somatic expansion for other repeat sequences and cell types would be required. MSH3 was identified as a modifier of X-linked dystonia-parkinsonism (XDP) onset, likely driven by SI of the CCCTCT repeat within the *TAF1* gene (*35, 36*). Interestingly, XDP and HD share neuropathological features, primarily the loss of MSNs in the striatum. The similarities with HD suggest AAV-DB-3.miMSH3 as potentially beneficial for XDP. MSH3 was also identified as a modifier of SI of the CTG repeat in the *DMPK* gene and disease severity in myotonic dystrophy type 1 (DM1) (*37, 38*). In summary, the integration of quantitative SI modeling with preclinical validation establishes a roadmap for developing and predicting mechanism-based disease-modifying therapies for HD and other repeat expansion diseases.

### Limitations of this study

The rate of somatic CAG expansion published by Handsaker et al. is based on a small number of patient donors. As the field matures, any updates to this rate can be integrated into the model framework. Because SI also occurs in cortical neurons (*5*), modeling cortical SI would allow for SI simulation across multiple brain regions and circuits affected in HD, affording more accurate predictions for widespread treatments. Finally, the HdhQ111 mouse model validates the human genetics studies through MSH3 lowering and impact on SI; however, it lacks robust functional phenotypes, such as mHTT aggregation and behavior deficits.

## STUDY DESIGN

The overall study design involved computationally simulating CAG repeat expansion in MSNs and predicting clinical benefit after targeting SI with a MSH3-lowering therapy. Preclinical proof-of-concept studies demonstrated AAV-mediated miMSH3 expression and lowering of MSH3 in NHPs and HD mice. AAV-mediated MSH3 lowering rates in NHP MSNs were used to predict clinical benefit across a broad range of HD patients. Biological MOA and impact of Msh3 lowering on somatic CAG repeat instability was evaluated in HD mice.

## MATERIALS AND METHODS

### Simulating CAG repeat expansion in MSNs

Our simulation of CAG repeat expansion was built in R and works by first instantiating a population of MSNs (plotted as dots) each with a CAG repeat as an attribute. Each CAG repeat begins the simulation at year 0 at the germline CAG length. With each successive year in a simulated patient lifetime the CAG repeat attribute of each cell is modified in concordance with the Handsaker two-phase linear model of CAG repeat expansion μ(x) = r1*max(x-T1,0) + r2*max(x-T2,0) (*2*). The stochastic nature of CAG repeat expansion leads to variable expansion that compounds with longer repeats expanding more rapidly and cells with shorter repeats more likely to persist at shorter lengths. As CAG repeats expand their length may surpass 150 CAGs and enter a period of continuous transcriptional dysregulation. We use this 150 CAG length as a threshold for classifying cells as healthy (<=150) and dysregulated (>150). We parameterized the rate of transduction and rate of SI lowering and here demonstrate the results of a simulated gene therapy.

### Predicting therapeutic benefit

To predict therapeutic benefit, we first run simulations as described above in both the presence and absence of a SI modifying therapeutic. Then we select an HD landmark for which we want to calculate therapeutic benefit. Example landmarks include the onset of HD-ISS stage (I,II, or III), or CAP-100. Then we identify the percentage of healthy cells remaining at the year the HD landmark of interest is reached in the untreated simulation. Then, by identifying the age at which the equivalent percentage of healthy cells is reached in the treated simulation, we calculated the difference in years between the untreated and treated simulations. This difference in age at which the HD landmark is reached is the predicted therapeutic benefit.

### AAV vector production

Recombinant AAV (rAAV) vectors were generated by the Research Vector Core at the Raymond G. Perelman Center for Cellular and Molecular Therapeutics at The Children’s Hospital of Philadelphia (CHOP). siSPOTR was used to design artificial miRNAs targeting human and NHP *MSH3* mRNA (miMSH3) (*27*). miMSH3s were embedded in the pri-miRNA miR-30 or miR-451 scaffold and cloned into a rAAV plasmid shuttle with AAV2 inverted terminal repeat sequences and kanamycin selection (*39*). rAAV vectors were produced by the standard calcium phosphate transfection method in HEK293 cells with the AdHelper plasmid, AAV-DB-3 Rep2/Cap1 peptide modified packaging plasmid, and rAAV shuttle plasmid with double CsCl purification (*23, 40*). Vector titers were determined by ddPCR.

### NHP procedures

All NHP procedures were conducted in accordance with the Guide for the Care and Use of Laboratory Animals (National Research Council) and were approved by the CHOP Research Institute Animal Care and Use Committee. Animals were housed under a 12-hour light:dark cycle with twice daily feedings with Purina LabDiet Certified Primate Diet (5048) enriched with fruits and vegetables and ad libitum access to purified drinking water.

NHPs received MRI-guided bilateral GP injections with AAV test articles or vehicle control as indicated in Table S1. AAV formulation buffer (CHOP Research Vector Core) was used as a vehicle control and to dilute AAVs to the target concentration prior to dosing. Animals were sedated with ketamine and xylazine and maintained on isoflurane anesthesia during the procedure. The ClearPoint Preclinical Orchestra and navigation software were used for targeting the GP. Injections were performed using a ClearPoint cannula (CUS-SMFL-03) attached to a syringe pump set to 1 µL/min. The two hemispheres were injected sequentially using the same cannula and an injection volume per hemisphere set to half of the total dose volume per animal in Table S1. A 10-minute dwell period was included following each injection before removing the cannula. Following both injections, the skin was sutured and the animal recovered, given buprenorphine SR as analgesia, and monitored postoperatively for pain and welfare.

After 4 weeks, NHPs were sedated and transcardially perfused with ice-cold saline. The brain was removed and sliced in 4-mm coronal slabs in a rhesus macaque brain matrix. For molecular analysis, brain regions of interest and other tissue types were collected and snap frozen in liquid nitrogen. For RNAscope FISH, brain slabs were postfixed in 10% neutral buffered formalin (NBF) for 24 hours, transferred to saline, and stored refrigerated for no more than 6 days before embedding in paraffin.

### RT-qPCR for gene expression

Total RNA was extracted from NHP brain samples using the Quick-DNA/RNA Miniprep Plus kit (Zymo Research D7003) and quantified with the Qubit BR Assay kit (Life Technologies Q10210). cDNA was generated from 500 ng total RNA with Maxima H minus reverse transcriptase (Thermo Scientific EP0752) and random hexamers (Life Technologies SO142). qPCR was performed with Luna Primer Probe 2X Master Mix (NEB M3004) including primer probe sets for rhesus *MSH3* (Applied Biosystems Rh00989001_m1) and *TBP* (Applied Biosystems Rh00427620_m1). Total RNA was extracted from HdhQ111 mouse brain samples with MagMAX mirVana total RNA isolation kit (Thermo Scientific A27828) and KingFisher Flex system using a qualified RT-qPCR method (BioAgilytix Labs, Durham, NC). Tissue samples were combined with 300 μL of lysis binding mix (containing lysis buffer and 2-mercaptoethanol). The samples were homogenized using TissueLyser II (Qiagen) with 5 mm stainless steel beads at 25 Hz for 2 x 3-minute cycles. The RNA eluates were quantitated using Qiagen QIAxpert. RT-qPCR was performed using the TaqMan Fast Virus 1-Step Master Mix (Applied Biosystems 4444432) and duplexed with primers and probes for mouse *Msh3* Mouse (Applied Biosystems Mm00487756_m1) and mouse *ActB ACTB* (Applied Biosystems Mm00607939_s1). Target gene Ct values were normalized to the housekeeping gene (ΔCt), then the average values were compared to average values of control samples (ΔΔCt). The 2^(-ΔΔCt)^ method was used to calculate fold change in gene expression relative to vehicle controls.

### Stem-loop RT-qPCR for miRNA expression

Total RNA was extracted from NHP brain samples using the Quick-DNA/RNA Miniprep Plus kit (Zymo D7003) according to the manufacture’s protocol. RNA samples were quantified by Qubit BR Assay kit according to the manufacturer’s protocol (Invitrogen Q10210). Stem-loop reverse transcription (RT) was carried out using TaqMan MicroRNA Reverse Transcription Kit, (Thermo Scientific 4366597 with 10 ng of RNA input and custom Taqman small RNA stem-loop primer (Life Technologies 4398988) targeting miMSH3. qPCR was carried out using Luna Primer Probe 2X Master Mix (NEB M3004). To quantify absolute miRNA levels, an HPLC purified RNA oligonucleotide (Integrated DNA Technologies) corresponding to the miMSH3 guide strand was used to generate a standard curve from a range of 1×10^2^ to 1×10^8^ copies per reaction to interpolate miMSH3 expression levels. Data were normalized to miRNA copies per ug of total RNA input.

### ddPCR for AAV biodistribution

Genomic DNA (gDNA) was extracted from NHP brain samples using the Quick-DNA/RNA Miniprep Plus kit (Zymo Research D7003) according to the manufacture’s protocol and quantified using Qubit dsDNA BR assay kit (Invitrogen Q32853). gDNA input for the GP, caudate, and putamen were 0.66 or 6.6 ng per well. Wells with saturated positive droplets were excluded. When more than one concentration was tested and concentrations were in range, copy numbers were averaged. ddPCR were performed on the QX200 (Bio-Rad) according to manufacturer’s instructions for probe-based assays with custom primer probe sets targeting the CAG promoter sequence (Integrated DNA Technologies) in the AAV genome and NHP *RPP30* sequence (Integrated DNA Technologies) as a reference for normalization. AAV vector genome copies were normalized to NHP *RPP30* copies to quantify AAV vector genomes per diploid genome using Bio-Rad QX Manager Software (v2.2.0.71).

### Capillary immunoassay (Jess)

Tissues were lysed in RIPA buffer (Pierce 89900) with 1X HALT protease inhibitor and homogenized using 5 mm stainless steel beads (Qiagen 69989) with Tissue Lyser LT at 50 Hz for 1 minute, incubated on ice for 30 seconds, and repeated for 3 cycles total, followed by incubation on ice for 5 minutes. Cell lysate debris was removed by centrifuging for 20 minutes at 16,000 g at 4 degrees Celsius, and supernatants were collected and stored in LoBind tubes at -80 degrees Celsius. Protein concentrations were quantified with Pierce BCA protein assay kit (ThermoFisher 23225). Samples were normalized to 2 mg/mL in 0.1X Sample Buffer and run on Jess capillary immunoassay (Protein Simple, SM-W001) following manufacturer’s protocol. Primary mouse anti-MSH3 (BD Biosciences 611390) and secondary anti-mouse HRP (Protein Simple DM-002) were used for detection of MSH3 using default parameters Data was analyzed using Compass for SW software (v6.3, Protein Simple). Data are plotted as MSH3 peak areas normalized to vehicle controls.

### RNAscope FISH

NHP brain slabs were trimmed, embedded in paraffin, and microtome-sectioned at a thickness of 5 microns at StageBio (Frederick, MD). Unstained tissue sections mounted on slides were shipped to CHOP and stored refrigerated with desiccants prior to staining. Fluorescence in situ hybridization (FISH) was performed using the RNAscope multiplex fluorescent reagent kit v2 assay (Advanced Cell Diagnostics, Cat. #323100-USM) following the manufacturer’s instructions. RNAscope probes *PPP1R1B* (DARPP-32; Cat. #1241971-C3) and *MSH3* (Cat. #1691821-C2) were used to detect mRNA transcripts in the putamen and caudate nucleus of rhesus macaques. Opal690 and Opal620 fluorophores (Akoya Biosciences, FP1497001KT and FP1495001KT) were used to detect the multiplexed *PPP1R1B*-C3 and *MSH3*-C2 probes, respectively. Slides were imaged using a Leica SP8 confocal microscope with LAS X v.3.7 software.

### Automated cell counting

Quantification of cells from fluorescence RNAscope FISH images was performed using Qupath. Cells were segmented using the “WatershedCellDetection” function. Nuclei were defined using the nuclear stain channel and, due to lack of a cytoplasmic stain, cytoplasmic regions were added using a 10 micron “cellExpansion” region around each nucleus. Subcellular *MSH3* mRNA puncta were quantified using the “SubcellularDetection” function. A custom groovy script was used to designate each subcellular detection as nuclear or cytoplasmic. A counts table was exported containing per cell puncta counts and additional measurements. Data management and visualization was performed using a custom R script.

### HdhQ111 mice

HdhQ111 KI heterozygous mice were previously described and bred at The Jackson Laboratory (Bar Harbor, ME) (*41*). Mice were shipped to PsychoGenics, Inc. (Paramus, NJ) for the in-life portion of the study. Procedures were approved by the Institutional Animal Care and Use Committee in accordance with the National Institute of Health Guide for the Care and Use of Laboratory Animals. Animals were balanced into treatment groups by body weight. At 8 weeks of age, test articles were administered to animals via simultaneous bilateral striatal injections. Standard aseptic surgical procedures were used to perform the injections. All injections were performed under isoflurane anesthesia (3-4% induction, 1-2% maintenance). The stereotaxic coordinates were 0.86 mm anterior and ±1.8 mm lateral with respect to bregma, and -2.5 mm dorsal ventral from the dura, and 5 µL volume per hemisphere was injected into the striatum at a 0.2 µL/minute infusion rate. Following infusion, the needles were left in place for 5 minutes to allow the test article to diffuse, the needle was then slowly retracted over 1-2 minutes. The skin was sutured over the injection site and the animal was returned to the home cage.

### Somatic instability index

Somatic instability testing was performed at Transnetyx (Culver City, CA). Genomic DNA was extracted using a magnetic bead-based extraction protocol. PCR amplification of the *Htt* gene containing CAG repeats was amplified using a Thermo Hot Start Master Mix and a set of primers, one of which was labeled with 6-FAM fluorescent dye. PCR amplification was conducted on an Simpliamp thermocycler using a touchdown PCR protocol to improve specificity and yield. Fragment analysis and sizing were performed by PCR products (2 µL) mixed with 0.5 µL of ROX1500 size standard and 7.5 µL of Hi-Di formamide. Samples were first normalized to ensure consistent input of 20 ng of DNA into the reaction. Samples were denatured at 96°C for 2 minutes and analyzed on an ABI 3730XL capillary electrophoresis platform. Data was processed using GeneMapper 5.0 software. CAG somatic instability index calculation was done using a Python script based on the method from Jong-Min Lee et al. (2010). Peak heights ≥1000 RFU were required for reliable instability index calculation. 10% thresholds were used for the striatum.

### Statistical analysis

Differences between untreated and treated groups were compared using one-way ANOVA with Dunnett’s test for multiple comparisons for mouse *Msh3* mRNA levels and somatic instability index. Differences between groups were considered to be significant at a *P* value of < 0.05. All results are shown as the mean ± standard deviation (SD). Statistical analyses were performed with GraphPad Prism v10.

## Acknowledgements

The authors acknowledge the Davidson lab and Dr. Ellie Carrell for thoughtful discussions, and Philip Morris for animal husbandry and planning. The authors thank Drs. Melanie McFadden and Alexis Mackiewicz and the entire Comparative Models Services Core and Department of Veterinary Resource staff at the Children’s Hospital of Philadelphia (CHOP). The authors thank Dr. Christina Torturo for conducting the in-life portion of the mouse pharmacology study at PsychoGenics, Inc.

## Funding

This work was supported by Latus Bio, the CHOP Research Institute, and the Hereditary Disease Foundation Transformative Research Award (BLD, JJC).

## Author contributions

Conception: BPS, PTR, DEL, GJY, PPG, JJC and BLD

Study design and execution: BPS, PTR, DEL, LT, RCG, IHO, ARH, NS, CMF, CPC, GJY, PPG, JJC and BLD

Writing & editing: BPS, PTR, DEL, GJY, PPG, JJC and BLD

## Competing interests

PTR, DEL and BLD are founders of Latus Bio. BPS, PTR, DEL, ARH, NS, CPC, CMF, JB, JCS, GJY, JC, PPG, and JJC are employees of Latus Bio. BLD has sponsored research and or serves an advisory role for Carbon Bio, Seamless Ther, and Latus Bio. PPG is a Latus Bio board member. All other authors declare that they have no competing interests.

## Data and materials availability

All data are available in the main text or the Supplementary Materials. All viruses and other materials may be available upon request to the corresponding authors through a material transfer agreement (MTA) from Latus Bio and/or CHOP Office of Technology Transfer. At the time of publication, a link to the source code will be provided.

