## Supplemental Figures and Table for "Computational modeling and preclinical validation support targeting somatic instability for Huntington’s disease treatment"

Bryan P. Simpson et al.

This file includes:

Figures S1 to S10

Table S1

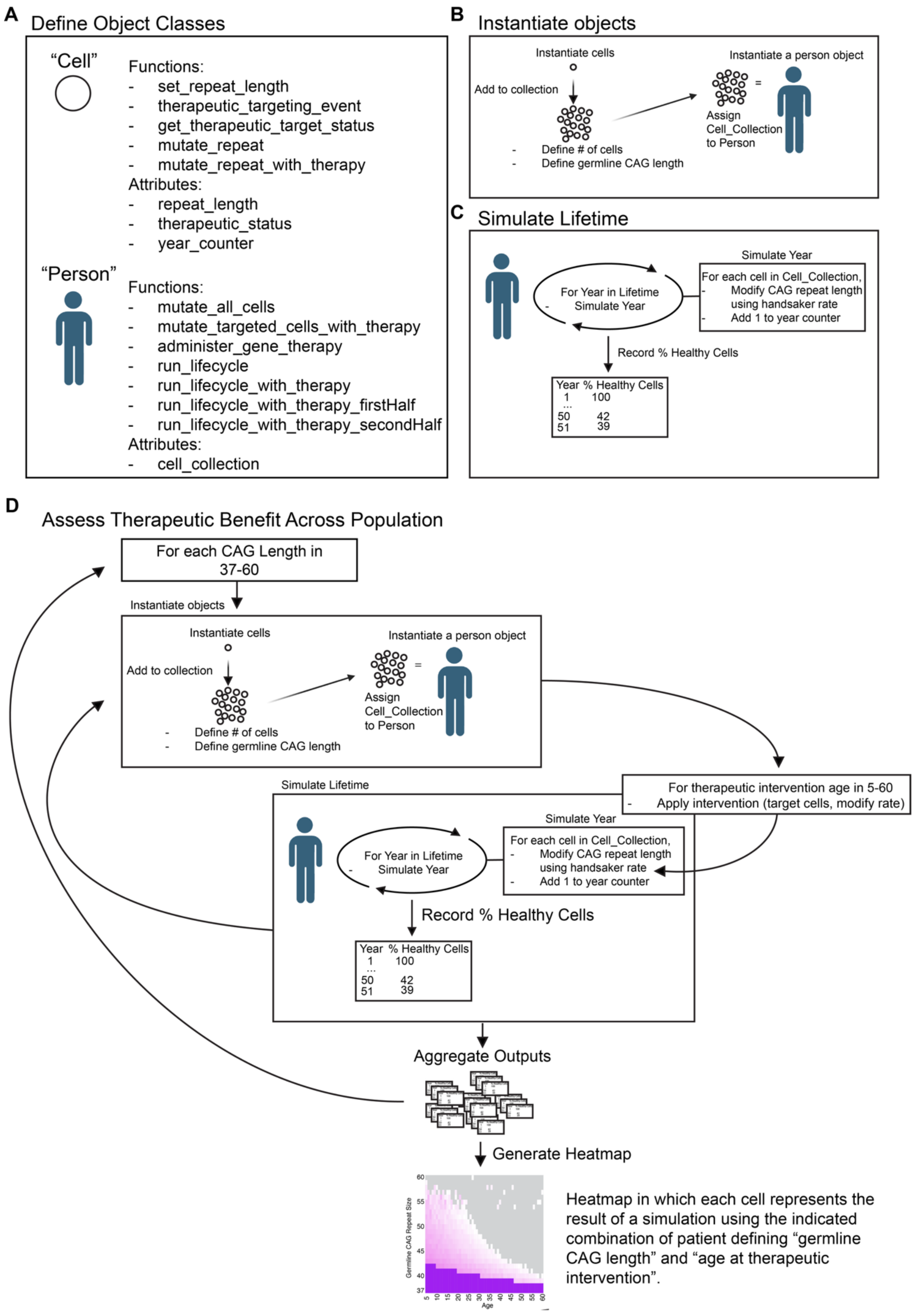

A
Define Object Classes
"Cell"
Functions:
- set_repeat_length
- therapeutic_targeting_event
- get_therapeutic_target_status
- mutate_repeat
- mutate_repeat_with_therapy
Attributes:
- repeat_length
- therapeutic_status
- year_counter
"Person"
Functions:
- mutate_all_cells
- mutate_targeted_cells_with_therapy
- administer_gene_therapy
- run_lifecycle
- run_lifecycle_with_therapy
- run_lifecycle_with_therapy_firstHalf
- run_lifecycle_with_therapy_secondHalf
Attributes:
- cell_collection
B
Instantiate objects
Instantiate cells
Add to collection
- Define # of cells
- Define germline CAG length
Instantiate a person object
Assign Cell_Collection to Person
C
Simulate Lifetime
For Year in Lifetime
- Simulate Year
Simulate Year
For each cell in Cell_Collection,
- Modify CAG repeat length using handsaker rate
- Add 1 to year counter
Record % Healthy Cells
Year % Healthy Cells
1 100
...
50 42
51 39
D
Assess Therapeutic Benefit Across Population
For each CAG Length in 37-60
Instantiate objects
Instantiate cells
Add to collection
- Define # of cells
- Define germline CAG length
Instantiate a person object
Assign Cell_Collection to Person
For therapeutic intervention age in 5-60
- Apply intervention (target cells, modify rate)
Simulate Lifetime
For Year in Lifetime
- Simulate Year
Simulate Year
For each cell in Cell_Collection,
- Modify CAG repeat length using handsaker rate
- Add 1 to year counter
Record % Healthy Cells
Year % Healthy Cells
1 100
...
50 42
51 39
Aggregate Outputs
Generate Heatmap
Germline CAG Repeat Size
60
55
50
45
40
37
Age
5
10
15
20
25
30
35
40
45
50
55
60
Heatmap in which each cell represents the result of a simulation using the indicated combination of patient defining "germline CAG length" and "age at therapeutic intervention".

**Figure S1: Simulation pipeline schematic. (A)** Object classes core to the simulation includes “cell” and “person.” Objects of class “cell” each contain attributes including a defined CAG length, a binary therapeutic status indicator, and a year counter. Cell objects can be modified by the indicated functions. Objects of class “person” contain a single attribute, a collection of simulated MSNs. By assigning all cells to a “person” object, a group of cells can be manipulated together across ages in a simulated lifetime. Objects of class “person” are modifiable by functions that apply mutation events to the CAG repeat at the prescribed rate. This rate may be modified by the simulated addition of an SI-targeted therapy in which cells are designated as targeted randomly in accordance with the provided rates of therapeutic efficacy (% of MSNs targeted and rate of SI lowering by % of MSH3 knockdown). **(B)** After class definition, cell objects must be instantiated. The number of created “cells” is user defined as is the germline CAG repeat length assigned to these cells. A “person” object is then instantiated and assigned a list of cell objects. **(C)** A lifetime is then simulated by modifying the CAG length each year in a user provided lifespan according to the Handsaker two-phase linear model of CAG repeat expansion. Simulation of therapeutic intervention can be integrated according to user provided rates of MSNs impacted and SI lowering. **(D)** SI-targeting therapies can be assessed across a population with ranges of germline CAG repeat lengths (e.g. 37-60) and ages at time of treatment (e.g. 5-60 years old) via a nested loop to iterate over both variable ranges, testing all provided CAG repeat lengths and ages at treatment combinations. Output measurements including therapeutic benefit (years of delayed onset) to a variety of HD landmarks (onset of HD-ISS Stage I, II, III, motor symptom onset and others) may be calculated this way.

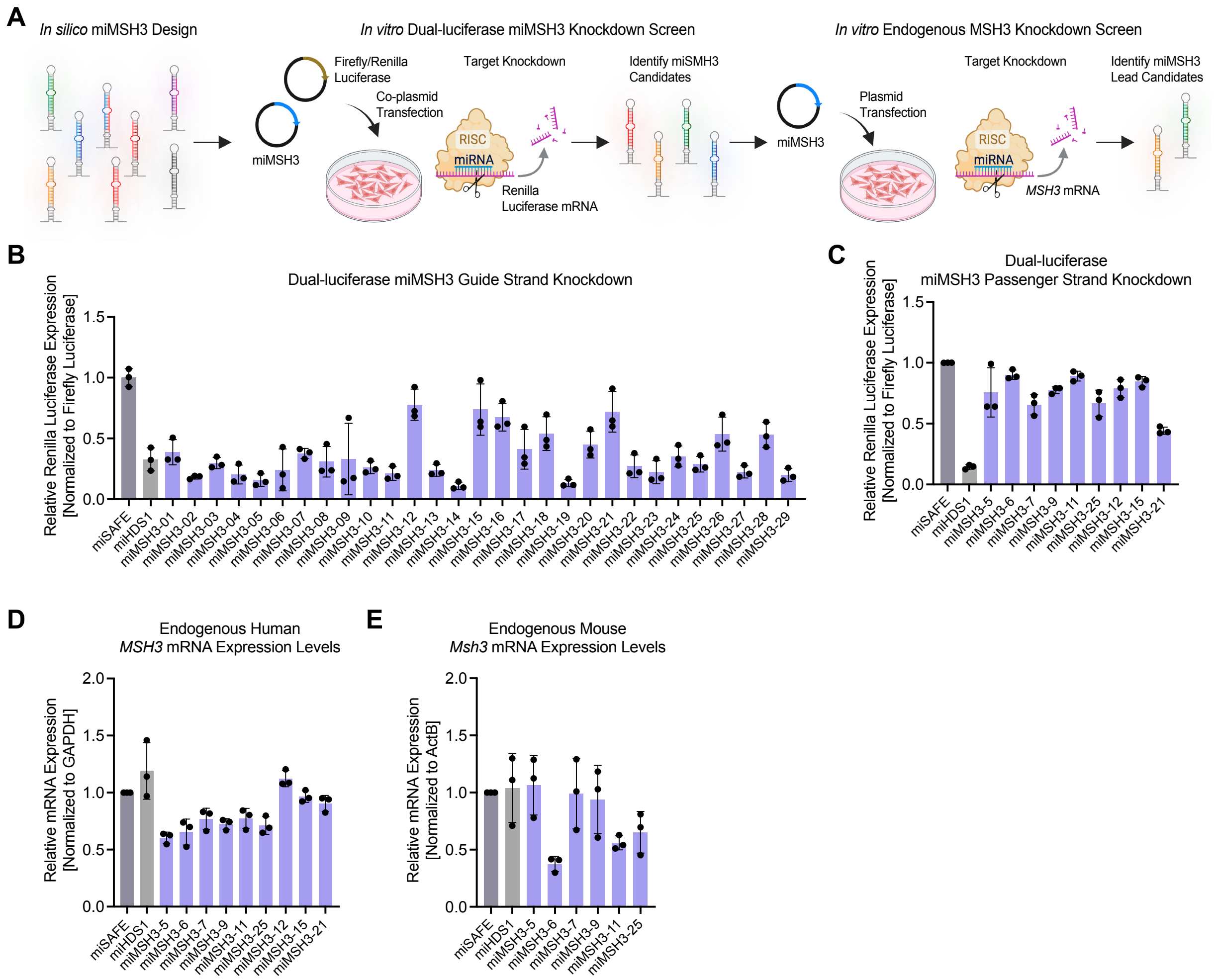


**Figure S2.** ***In vitro*** **screening identifies potent artificial miRNAs targeting MSH3. (A)** Schematic depicting MSH3 miRNA (miMSH3) *in silico* design with siSPOTR (26) and *in vitro* miMSH3 candidate screening workflow. **(B)** Twenty-nine miMSH3 guide strand sequences embedded in the miR-30 scaffold were screened by dual-luciferase knockdown assay in HEK293 cells after co-plasmid transfection of Renilla luciferase/firefly luciferase and miMSH3 expression plasmids. The miHDS1 (HTT-targeting miRNA) guide strand target was included as a knockdown positive control for the assay. Renilla luciferase (guide stand target plasmid) expression was normalized to firefly luciferase expression. **(C)** Nine miRNA candidates were screened for passenger strand bias and knockdown by dual-luciferase assay. **(D)** Nine miMSH3 sequences were screened for knockdown of endogenous human *MSH3* mRNA normalized to *GAPDH* in HEK293 cells by RT-qPCR. **(E)** Six miMSH3 sequences were screened for knockdown of endogenous mouse *Msh3* mRNA normalized to *ActB* in mouse N2a cells by RT-qPCR. N=3 independent experiments with three technical replicates, data are shown as the mean, and error bars represent standard deviation (SD). Panel A was created using BioRender.com.

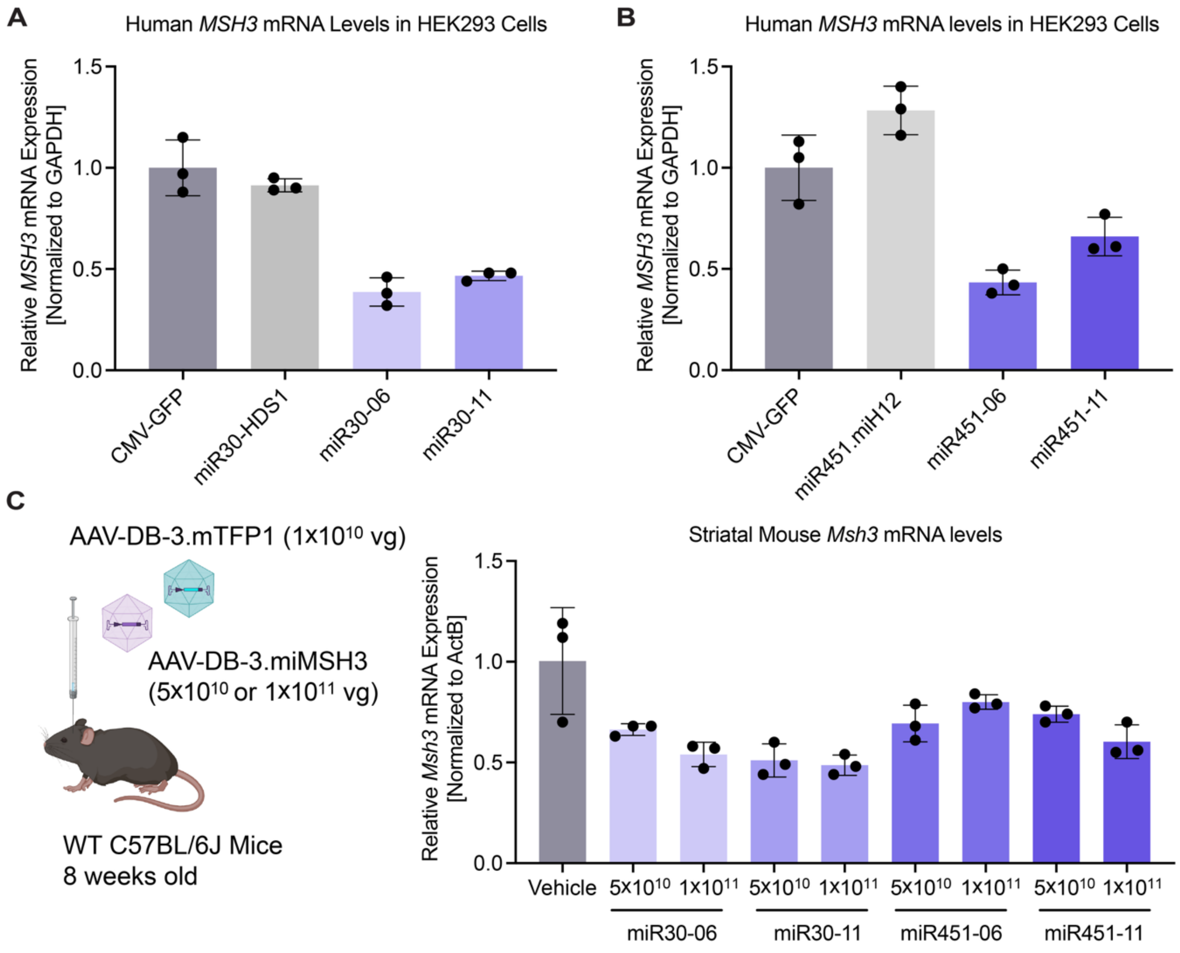


**Figure S3. miR-30 and miR-451 as scaffolds for miMSH3 mediated MSH3 reduction. (A)** Human *MSH3* mRNA reduction with miRNA miR-30 scaffold relative to no miRNA control in HEK293 cells by RT-qPCR. miR-30.miHDS1 targeting *HTT* mRNA was included as positive control. **(B)** Human *MSH3* mRNA levels after miRNAs with miR-451 scaffold. miHD12 targeting *HTT* was included as positive control. For (A) and (B), GFP-positive cells were sorted by FACS prior to RT-qPCR. N=3 independent experiments with three technical replicates and error bars represent SD. **(C)** C57BL/6J wildtype mice were bilaterally infused in the striatum at 8 weeks old with AAV-DB-3-miR-30.miMSH3 or AAV-DB-3.miR-451.miMSH3 at $5\times10^{10}$ vg or $1\times10^{11}$ vg with co-delivery of AAV-DB3-mTFP1 at $1\times10^{10}$ vg to guide microdissection of transduced tissue at 4 weeks post-infusion. Mouse *Msh3* mRNA levels were measured by RT-qPCR and were normalized to *ActB* and plotted relative to vehicle control. N=3 mice per group shown as mean and error bars represent SD. Panel C was created using BioRender.com.

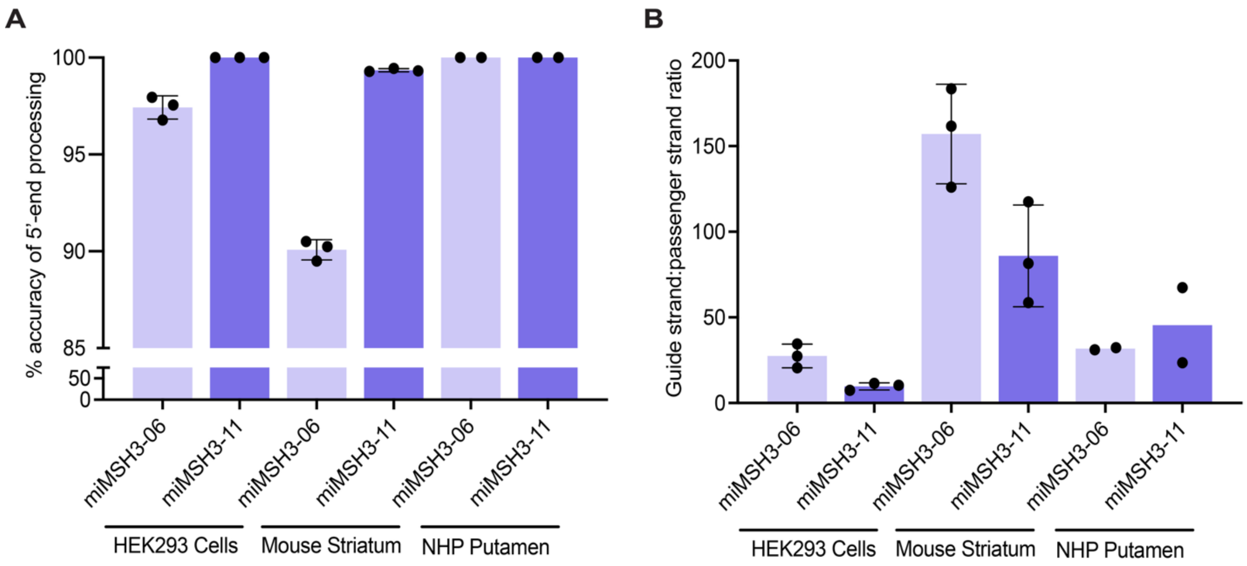


**Figure S4. Favorable miRNA processing in HEK293 cells, mouse striatum, and NHP putamen. (A)** The percent miR-30 mature guide strands with accurate 5'-end processing in the seed sequence, P2-P8, following expression in HEK293 cells, mouse striatum and NHP putamen as assessed by small RNA sequencing. **(B)** Guide to passenger strand ratios of miMSH3-06 and miMSH3-11 in HEK293 cells, mouse striatum and NHP putamen, using small RNA sequencing. N=3 independent experiments in HEK293 cells, N=3 WT mice, N=2 NHPs. Data are shown as the mean and error bars represent SD.

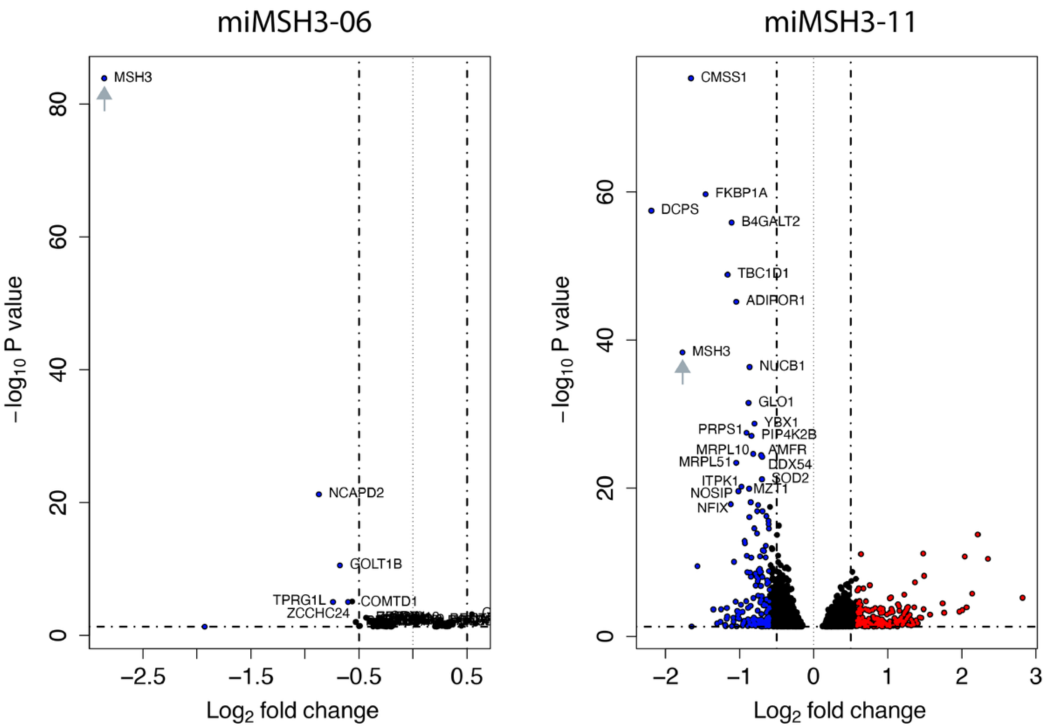


**Figure S5**. **Low detectable off-target gene expression in HEK293 cells transfected with miMSH3-06.** Volcano plots depict off-target differential gene expression determined by RNA sequencing of miMSH3-06 and miMSH3-11 transfected HEK293 cells, relative to a GFP reporter plasmid control. Data were cleaned using miHDS1 (*HTT*-targeting miRNA) control to remove shared transcriptional changes from plasmid transfection. Volcano plots indicate significance of observed differential gene expression on the x-axis (Log2 fold change) and amplitude of up (red dots) and down (blue dots) regulated gene expression on the y-axis (-log10 P value). Dotted lines indicate the thresholds for significance; non-significant differentially expressed genes are shown as black dots. Grey arrows indicate *MSH3*.

A

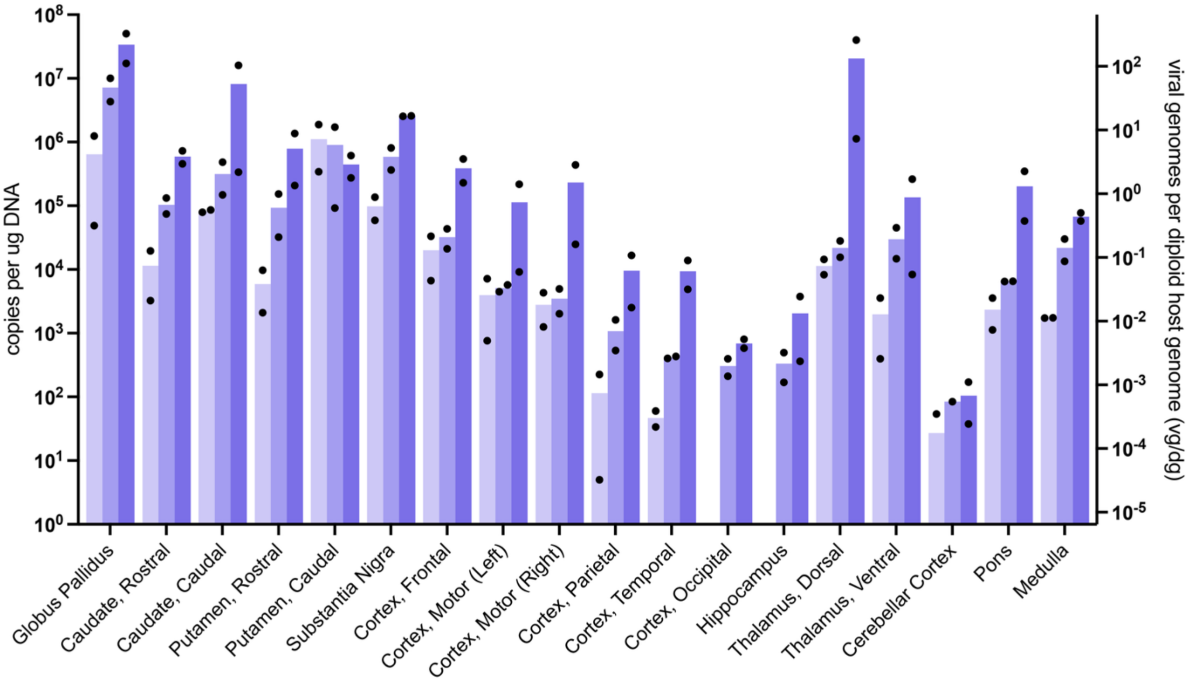


B

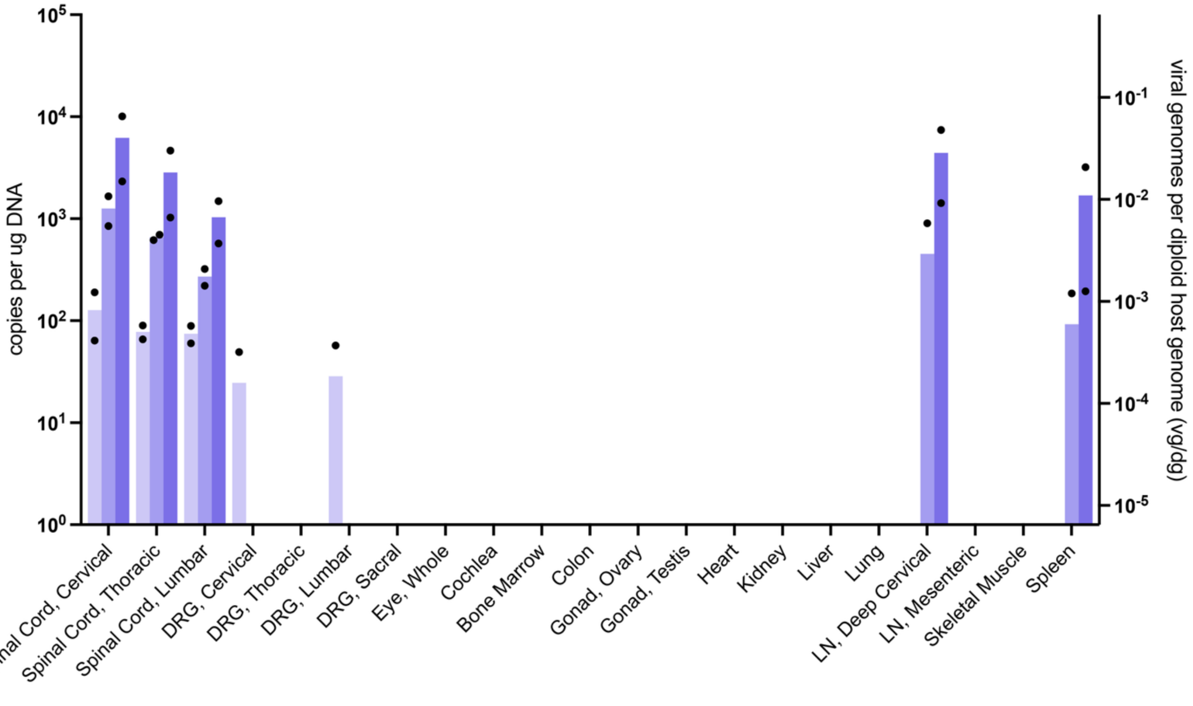


**Figure S6. High AAV biodistribution in NHP brain with minimal detection in other tissues.** AAV vector genome biodistribution data by qPCR are shown as copies per microgram of DNA (left y-axis) and calculated AAV vector genomes (vgs) per diploid host genome (dg) (right y-axis) across brain tissues **(A)** and other tissues **(B)**. Dose groups were $1.5{\times}10^{11}$ vg, $8.2{\times}10^{11}$ vg, and $3.2{\times}10^{12}$ vg shown as light purple, purple, and dark purple bars, respectively. Biodistribution data not shown were below the limit of quantification (BLOQ). Data are shown as group mean (N=2) with data points from each animal. Samples from vehicle-treated control NHPs were below the limit of detection (BLOD). In plots, group means were calculated using a value of 0 for any result below the limit of quantification (BLOQ).

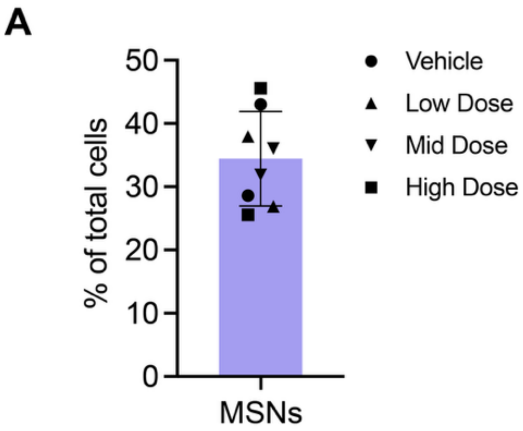


Total MSN Count: 1761

Total Cell Count: 5179

**Figure S7. MSNs represent 34.4% of total cells in the rhesus macaque striatum. (A)** RNAscope FISH counts of *PPP1R1B+* MSNs and total cells counterstained with Hoechst were used to calculate the percentage of total cells that are MSNs in the striatum (caudate and putamen) of rhesus macaques. Data are shown as the mean from the vehicle (circles), $1.5\times10^{11}$ vg low dose (upward triangles), $8.2\times10^{11}$ vg mid dose (downward triangles), and $3.2\times10^{12}$ vg high dose (squares) groups (N=2 NHPs/group) and the error bar represents SD. The total counts for MSNs and all cells profiled is shown below the plot.

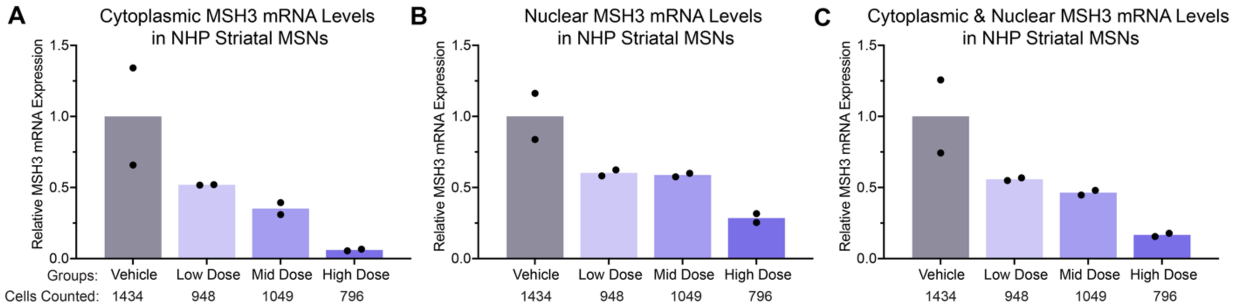


**Figure S8.** ***MSH3*** **mRNA lowering in striatal MSNs. (A)** RNAscope FISH quantification of cytoplasmic *MSH3* mRNA puncta counts in all *PPP1R1B+* MSNs profiled from the striatum were determined with an automated cell segmentation and puncta counting algorithm. **(B)** RNAscope quantification of nuclear *MSH3* mRNA puncta counts in all *PPP1R1B+* MSNs profiled from the striatum. **(C)** RNAscope quantification of combined cytoplasmic and nuclear *MSH3* mRNA puncta counts in all *PPP1R1B+* MSNs profiled from the striatum. Data plotted relative to vehicle control group and shown as group mean for the $1.5{\times}10^{11}$ vg low dose, $8.2{\times}10^{11}$ vg mid dose, and $3.2{\times}10^{12}$ vg high dose groups (N=2 NHPs/group) with data points that represent each animal.

**A**

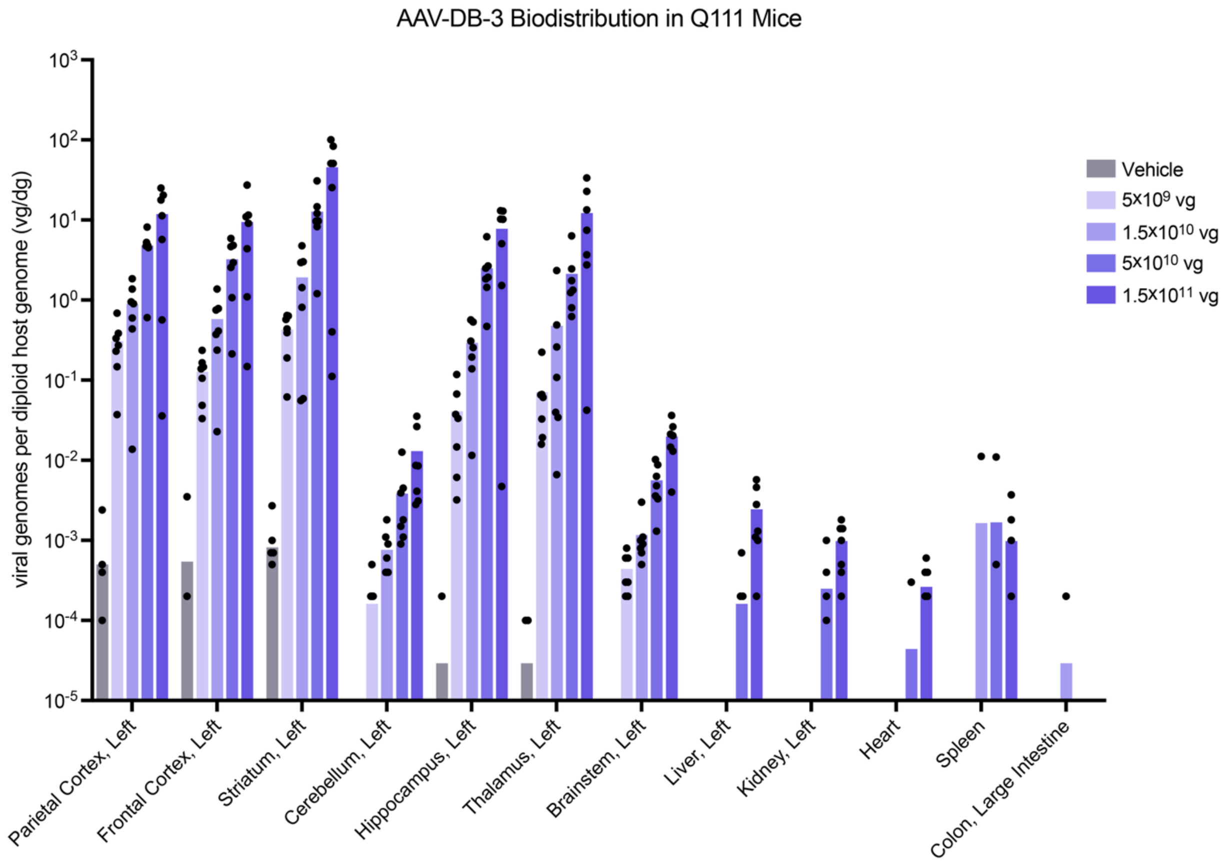


**Figure S9. Dose-dependent AAV-DB-3 biodistribution in HdhQ111 mouse brain tissues with minimal detection in peripheral tissues. (A)** AAV vector biodistribution data by qPCR are shown as calculated AAV vector genomes (vgs) per diploid host genome (dg) (y-axis) across brain tissues and other tissues. Biodistribution not shown were below the limit of quantification (BLOQ). Data are shown as group mean (N=7, vehicle group; N=7, $5\times10^{9}$ vg group; N=7, $1.5\times10^{10}$ vg group; N=7, $5\times10^{10}$ vg; N=7, $1.5\times10^{11}$ vg group) with data points from each animal. In plots, group means were calculated using a value of 0 for any result below the limit of quantification (BLOQ).

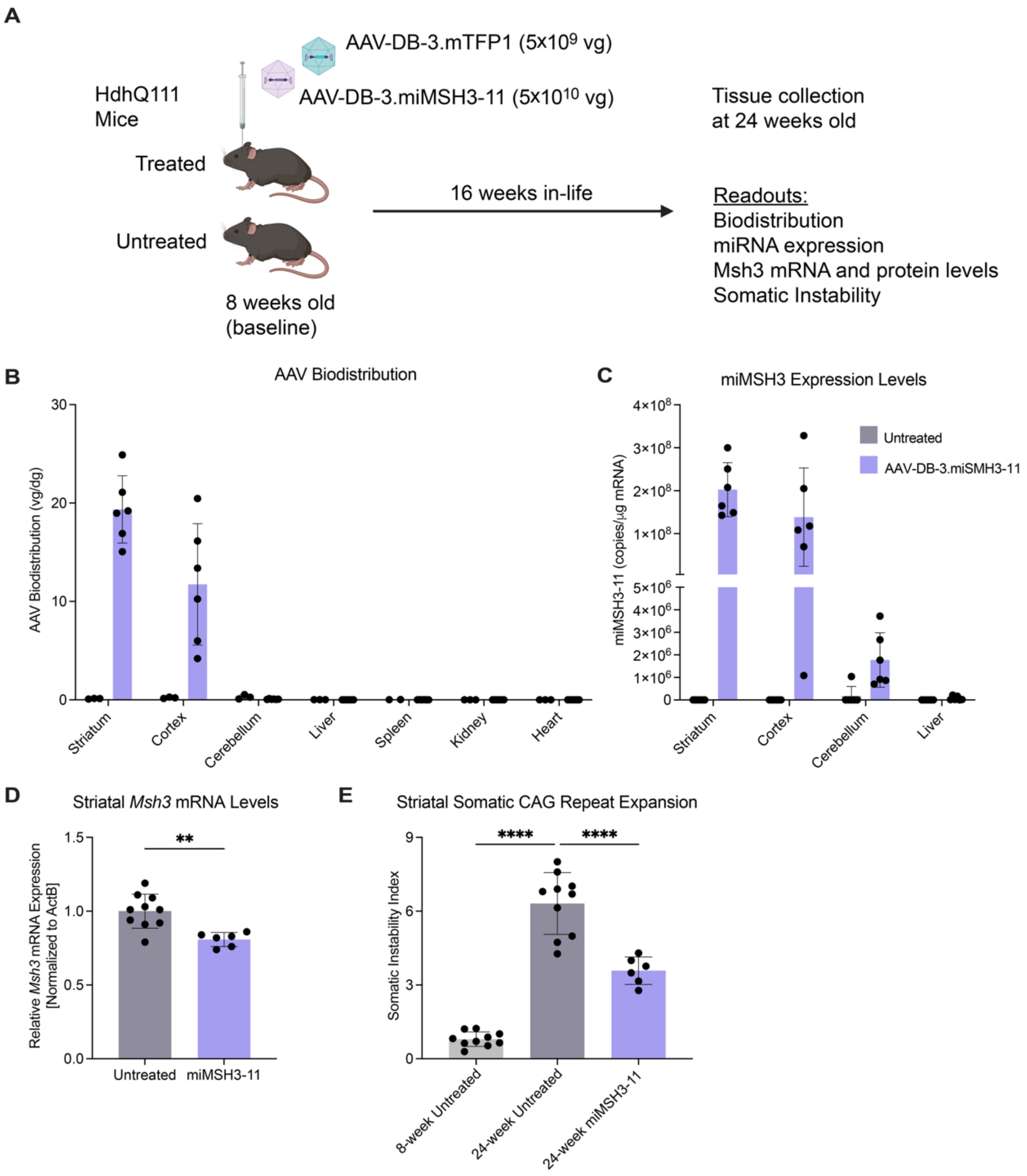


**Figure S10. AAV-DB-3.miMSH3-11 reduces somatic instability in HdhQ111 mice. (A)** Study design for evaluating intrastriatal infusion of AAV-DB-3.miMSH3-11 at $5\times10^{10}$ vg in heterozygous HdhQ111 KI mice. Co-infusion of AAV-DB-3.mTFP1 at $5\times10^{9}$ vg was performed to enable microdissection of transduced mTFP1-positive striatal tissue. **(B)** AAV vector biodistribution by ddPCR in untreated and treated HdhQ111 mice at 24 weeks old (N=3, untreated; N=6, AAV-DB-3.miMSH3-11 treated). **(C)** miMSH3-11 expression levels as measured by stem-loop RT-qPCR in untreated and treated HdhQ111 mice at 24 weeks old (N=6, untreated; N=6, AAV-DB-3.miMSH3-11 treated). **(D)** *Msh3* mRNA levels in the striatum in untreated and treated HdhQ111 mice at 24 weeks old by RT-qPCR. *Msh3* mRNA was normalized to *ActB* and plotted relative to untreated controls (N=10 untreated, N=6 treated; unpaired t-test, $**p < 0.01$). **(E)** SI index in untreated and treated mice by PCR-based fragment analysis (N=10, baseline untreated 8-week-old mice; N=10, untreated 24-week-old mice; N=6, AAV-DB-3.miMSH3-11 treated 24-week-old mice; one-way ANOVA with Dunnett's post hoc analysis, $****p < 0.0001$). Data are shown as the mean and error bars represent SD. Panel A was created using BioRender.com.

**Table S1: Summary of Study Design for NHP DRF Study**

| NHP # | Species | Sex | Test Article(s) | Total Dose (vg/NHP) | Total Dose Volume (µL/NHP) | Site of Administration | Associated Figure(s) |
|---|---|---|---|---|---|---|---|
| 1 | Rhesus macaque | M | Vehicle Control | 0 | 120 | Bilateral GP | Fig 4, 5, S7 |
| 2 | Rhesus macaque | F | Vehicle Control | 0 | 60 | Bilateral GP | Fig 4, 5, S7 |
| 3 | Rhesus macaque | M | AAV-DB-3. miMSH3-06 | $1.5\times10^{11}$ | 60 | Bilateral GP | Fig 4, 5, S4, S6, S7 |
| 4 | Rhesus macaque | F | AAV-DB-3. miMSH3-06 | $1.5\times10^{11}$ | 60 | Bilateral GP | Fig 4, 5, S4, S6, S7 |
| 5 | Rhesus macaque | M | AAV-DB-3. miMSH3-06 | $8.2\times10^{11}$ | 120 | Bilateral GP | Fig 4, 5, S6, S7 |
| 6 | Rhesus macaque | F | AAV-DB-3. miMSH3-06 | $8.2\times10^{11}$ | 120 | Bilateral GP | Fig 4, 5, S6, S7 |
| 7 | Rhesus macaque | M | AAV-DB-3. miMSH3-06 | $3.2\times10^{12}$ | 120 | Bilateral GP | Fig 4, 5, S6, S7 |
| 8 | Rhesus macaque | F | AAV-DB-3. miMSH3-06 | $3.2\times10^{12}$ | 120 | Bilateral GP | Fig 4, 5, S6, S7 |
| 9 | Rhesus macaque | F | AAV-DB-3. miMSH3-11 | $1.5\times10^{11}$ | 60 | Bilateral GP | Fig S4 |
| 10 | Rhesus macaque | F | AAV-DB-3. miMSH3-11 | $1.5\times10^{11}$ | 60 | Bilateral GP | Fig S4 |
